# NPMc+ and Flt3-ITD cooperation promotes a new oncogenic HSC state that supports transformation and leukemia stem cell maintenance

**DOI:** 10.1101/2024.07.25.605115

**Authors:** Maria Elena Boggio Merlo, Maria Mallardo, Lucilla Luzi, Giulia De Conti, Chiara Caprioli, Roman Hillje, Mario Faretta, Cecilia Restelli, Andrea Polazzi, Valentina Tabanelli, Angelica Calleri, Stefano Pileri, Pier Giuseppe Pelicci, Emanuela Colombo

## Abstract

*NPMc+* and *FLT3-ITD* are the most frequently co-occurring mutations in Acute Myeloid Leukemia (AML). Cellular and molecular mechanisms of their cooperation are largely unknown. We investigated their effects on pre-leukemic and leukemic hematopoietic stem cells (HSCs) in mouse models that recapitulate the human disease. Both oncogenes increase proliferation of pre-leukemia HSCs, however, only *NPMc+* extends self-renewal, by preventing depletion of quiescent HSCs. Quiescent HSCs include dormant and active HSCs, which are dynamic and alternative phenotypic states supporting, respectively, self-renewal and regenerative haematopoiesis. Analyses of the dormant and active transcriptional programs showed that NPMc+ stimulates the dormant-to-active transition but also enforces dormancy, thus allowing dormant HSCs to be replenished. Co-expression of NPMc+ and *Flt3-ITD* induces a new phenotypic state of quiescent HSCs whereby dormancy and activity co-exist at single-cell level. This new state likely supports *in vivo* proliferation of self-renewing HSCs and rapid selection of leukemia initiating cells. Pharmacological inhibition of the TGFb cell-dormancy pathway reduces self-renewal of leukemia SCs and prolongs mouse survival allowing to conclude that enforcement of HSC dormancy is a critical determinant of unrestricted self-renewal during leukemia development and represents a target for novel anti-leukemic strategies.

## Introduction

Progression towards Acute Myeloid Leukemia (AML) is driven by the selection of multiple genetic mutations (two to six recurrent mutations), which confer different adaptive phenotypes to the target cells (clonal evolution) and manifest clinically with progressively-increasing disease severity(1).

Mutations in the Nucleophosmin gene (*NPMc+*) are the most frequent genetic alterations in AMLs (about 30%) and are often found within the same patient together with mutated *FLT3 (FLT3-ITD*) and/or *DNMT3A* (*DNMT3Amut*) suggesting cooperation among these mutations (2), as confirmed by transgenic mice expressing doubles or triple mutants (3–7).

In patients, the prognosis of the NPM1c+/FLT3-ITD double-mutant is still very poor and new therapeutic strategies are urgently needed for these AMLs. However, the relative contribution of *NPMc+* and *FLT3-ITD* to the development of AML is unclear, as well the cellular and molecular bases of their cooperation.

A specific pathway of the AML clonal-evolution has been characterized, based on the identification of somatic mutations in the peripheral blood of healthy people (Clonal Hematopoiesis or CHIP) in genes (*DNMT3A, TET2, JAK2, ASXL1, SF3B1*) that are frequently mutated also in indolent cytopenia (CCUS), myelodysplastic syndrome (MDS), myeloproliferative neoplasms (MPNs) or AMLs (8, 9).

In AMLs with CHIP carrying mutations of both *NPM*1 and *DNMT3A*, *NPMc+* is never found in T-lymphocytes (0 of 12 cases analysed)(10). Analyses of non-leukemic HSC/progenitors and mature populations, however, showed the presence of *DNMT3Amut* across different populations in all cases, and of *NPMc+* selectively in myelo-lymphoid and/or granulocyte-monocyte progenitors in ∼50% of cases (6 of 11), suggesting that *DNMT3A* mutations arise in HSCs early in AML evolution, while *NPM* mutations may occasionally arise in the pre-leukemic myeloid progenitors (10). Expression of *Npmc+* in mouse myeloid progenitors, alone or in the presence of *Dnmt3Amut*, induces extended self-renewal and HSC-reprogramming, thus establishing a HSC/progenitor pre-leukemic population that eventually evolves into frank leukemia (11). Thus, *NPM* mutations may drive clonal evolution of progenitors in the pre-leukemic CHIP bone-marrow. The majority of AMLs with double *NPM1*-*DNMT3A* mutations, however, do not show evidence of CHIP (∼60%;(12)), suggesting that *NPMc+* may play different functions during AML development.

*NPM* mutations typically occur in *de novo* AMLs. They are usually found in the dominant leukemic clone together with other sub-clonal mutations, such as *FLT3-ITD* (2, 13) and are present at relapse in ∼90% of patients(14), suggesting that they are critical for *t*he appearance of the fully-transformed leukemia phenotype (AML-founding mutations). However, though AML with *NPM1* mutation is a distinct genetic entity in the revised WHO classification, patients with *NPMc+* are heterogenous with respect to patterns of co-mutation, prognosis and therapy (15). Analyses of transcription patterns across a large cohort of *NPMc+* AMLs identified two major subtypes, classified as either “committed” or “primitive”. The “committed” subtype has better survival and is enriched in *DNMT3A* mutations, myeloid-differentiation genes and cells with aberrant myelo-monocytic differentiation, while the primitive subtype has worse survival and is enriched in *FLT3-ITD* mutations, stem/progenitor genes and stem/progenitor-like cells. Notably, NPMc+ has been documented in HSC-enriched CD34+ cells (16), as well in myeloid, monocytic, erythroid and megakaryocytic cells in the bone marrow of NPMc+AML patients (17). Moreover, the only reported case of *de novo NPM1c+/DNMT3A_WT* AML for which the pre-leukemic bone marrow has been investigated, clearly showed occurrence of the *NPM1* mutation in the pre-leukemic HSCs (18). Together, these data suggest that *NPMc+* might target different cell compartments (progenitors or HSCs), depending on the trajectories of clonal development (CHIP or *de novo* AMLs), or types of co-mutations (*DNMT3A or FLT3-ITD*). Considering the relevance that the developmental biology of the disease has in the design of an effective treatment(19), recent studies have addressed the impact of different AML associated mutation on the biology of the hematopoietic cells in the pre-leukemic phase of the disease in the attempt to define underline oncogenic mechanism that could lead to the development of new therapeutic interventions(20, 21). We report here that enforcement of quiescence/dormancy by NPMc+ is critical for the establishment of extended self-renewal in pre-leukemic HSCs, and we further investigated its role in *NPMc+/FLT3-ITD* cooperation in AML initiation and growth.

## Results

### NPMc+ increases HSC proliferation and self-renewal

To investigate the effects of NPMc+ on HSCs we used transgenic mice carrying a conditional loxP-flanked allele of the NPMc+ human cDNA inserted in the *Hprt* locus(3). Upon conditional expression of the transgene, mice showed increased expression of *Hox*-genes in the bone marrow (BM) lineage negative (lin-) compartment, and leukemia development at low penetrance and long latency(3), as reported for the other available NPMc+ mouse model where mutated NPM1 is expressed under the control of the *Npm1* endogenous promoter (4). To follow expression of the NPMc+ transgene *in vivo*, NPMc+ mice were crossed with a syngeneic strain carrying a loxP-flanked Rosa26-EYGF allele(22) (thereafter NPMc+/YFP and YFP respectively). BM mononucleated cells (BM-MNCs) isolated from NPMc+/YFP or control YFP mice were treated *in vitro* with the recombinant TAT-CRE protein and FACS-sorted to separate YFP+ and YFP-subpopulations. As shown in Fig. 1A, NPMc+ expressing cells are present only in the YFP+ population of NPMc+/YFP mice, with ∼85% of the YFP+ population also expressing NPMc+ (Fig. 1B).

**Figure 1.**
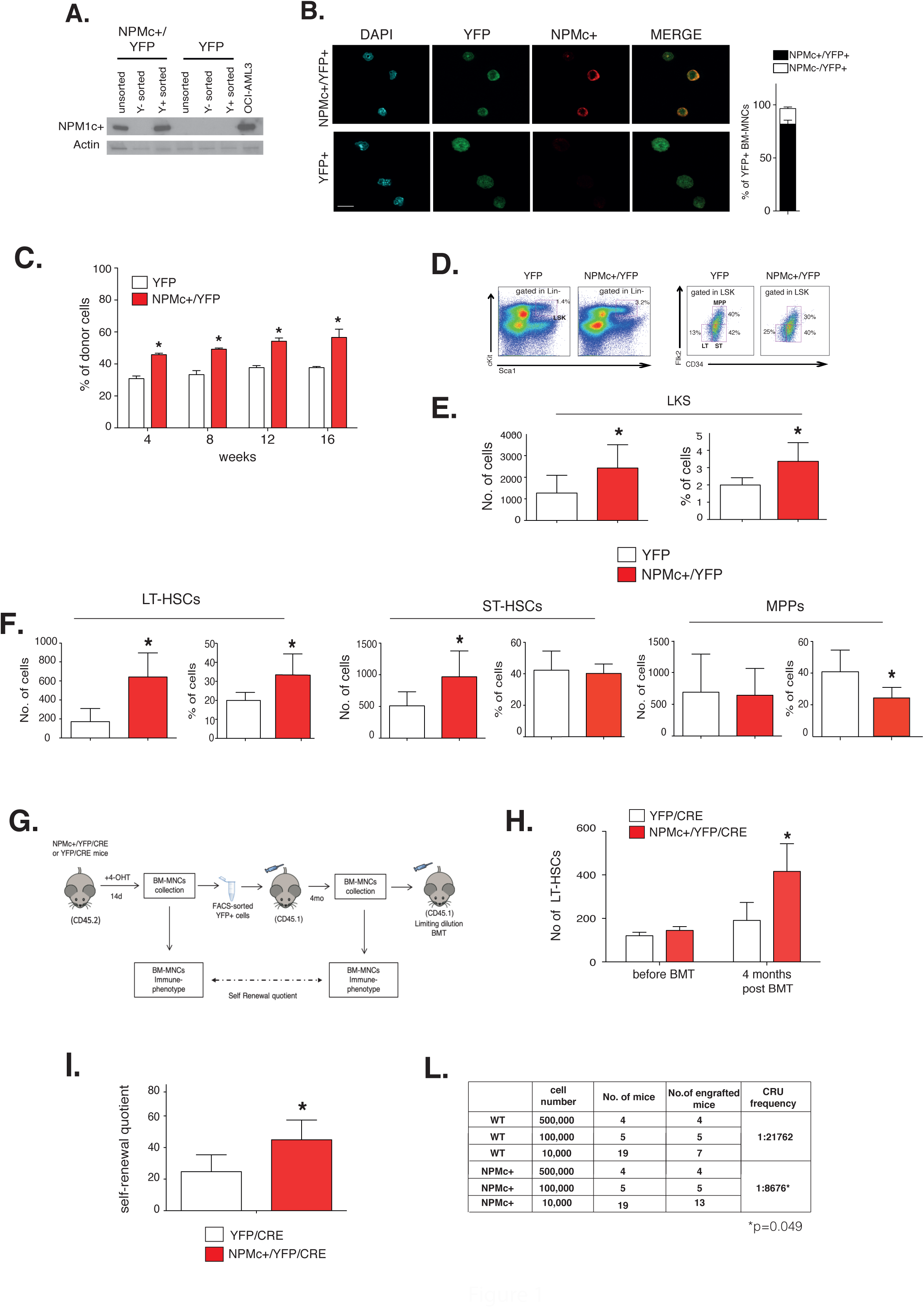
NPM1c+ expression promotes preleukemic HSC expansion and self-renewal. **A.** Western Blot analysis of NPMc+ expression in the YFP+ (Y+) and YFP- (Y-) fractions after FACS sorting of BM-MNCs isolated from NPMc+/YFP and YFP mice and treated *in vitro* with the TAT-CRE. Lysates from the OCI-AML3 NPMc+ cell line were used as positive control. Actin was used as a loading control. **B.** Left: Representative confocal images of anti-NPMc+ immunofluorescence analyses performed on YFP+ cells sorted from NPMc+/YFP and YFP mice: blue – DAPI; green – YFP; red – NPMc+ (original magnification 256X, scale bar 10μm). Right: percentage of NPMc+ positive cells within the YFP+ sorted population. **C.** Competitive BMT (1:1 ratio): percentage of CD45.2+ donor derived YFP+ cells in the PB of transplanted mice at the indicated time points (3-5 mice per cohort, graph representative of 1 of 3 independent experiments; *=p<0.05). **D.** Representative FACS gating schemes for the analysis of different hematopoietic populations in the BM of YFP and NPMc+/YFP animals. left: gating of the LSK (c-Kit+, Sca-1+, Lin-) population within lineage negative cells; right: gating of LT-HSCs (Lin-, Sca-1+, cKit+, CD34-, FLK-), ST-HSCs (Lin-, Sca1+, cKit+, CD34+, FLK-), and MPPs (Lin-, Sca1+, cKit+, CD34+, FLK+) within LKS cells. **E.F.** Quantification of the subpopulation depicted in D: numbers (per million of BM-MNCs, left panels) and percentages (right panels) of LSK in the Lin- population (E) and LT-HSC, ST-HSC and MPP in the LSK population (F), determined by FACS analyses in NPMc+/YFP and YFP BM-MNCs (3-5 mice per cohort; graph representative of 1 of 4 independent experiments; *=p<0.05). **G.** Self-Renewal assay and limiting number BMT experimental scheme: YFP/CRE and NPMc+/CRE/YFP recombined BM-MNCs have been FACS-analysed, sorted and transplanted in recipient mice. After 4 months, the BM has been collected, FACS-analysed and re-transplanted in a limiting dilution assay. **H.** Number of LT-HSCs in one million BM-MNCs derived from 4-OHT treated NPMc+/YFP and YFP mice, before (Input) and 4 months after BMT (5 mice per cohort, graph representative of 1 of 2 independent experiments; *=p<0.05). **I.** Self-renewal quotient calculated as the ratio between the number of donor LT-HSCs in the total BM of recipient mice at 4 months post BMT, and the numbers of transplanted LT-HSCs (*=p<0.05). **L.** Limiting BMT assay using different amount of YFP+ FACS-sorted BM-MNCs from NPM1c+/YFP or YFP mice (4 months after the first BMT). Engrafted animals are defined as recipients with >0.1% donor-derived PB cells, 4 months post BMT. The frequency of functional HSCs (Competitive Repopulation Units, CRUs) was calculated with the ELDA software. Mean ± s.d. values are shown. Unpaired Student’s t test has been applied and significant p values are reported.

We first examined the repopulating capacity of the pre-leukemic NPMc+ BM by competitive congenic BM transplantation (BMT). Equal numbers of FACS-sorted YFP+ BM-MNCs, purified from NPMc+/YFP or YFP (CD45.2^+^) and treated with TAT-CRE *in vitro,* were co-transplanted into lethally irradiated congenic (CD45.1^+^) mice along with CD45.1 BM-MNCs (1:1 ratio). The repopulating ability of donor cells was evaluated at different time points after transplantation (4 to 16 weeks) by analyzing the percentage of donor CD45.2+ white blood cells (WBC) in the peripheral blood (PB) of recipient mice. Mice reconstituted with NPMc+/YFP cells showed, at all time-points, significantly higher proportions of donor WBCs (Fig. 1C), demonstrating that NPMc+ confers higher BM repopulating potential.

To investigate whether the increased repopulating potential of the NPMc+_BM-MNCs correlates with the expansion of NPMc+ HSCs and/or early progenitors, we analyzed cell composition and cell cycle properties of the BM LSK cell population (Lin^-^, Sca^+^, c-Kit^+^), which is composed of Long Term-HSCs (LT-HSCs; CD34^-^ and FLK3^-^), Short Term-HSCs (ST-HSCs; CD34^+^ and FLK3^-^) and Multi-Potent Progenitors (MPPs; CD34^+^ and FLK3^+^). Mice were transplanted with TAT-CRE-treated NPMc+/YFP+ or YFP+ BM-MNCs and, four months after bone marrow transplantation (BMT), they were injected with a short-pulse of Bromodeoxyuridine (BrdU; 2 injections in 12 hours) prior to BM recovery. YFP+ BM-MNCs were then analyzed by FACS using lineage-specific(23) and anti-BrdU antibodies (FACS-analysis gating strategy for each sub-population is shown in Fig. 1D). The NPMc+/YFP+ bone marrow cells showed a significant expansion, in terms of both numbers and percentages, of LSK, LT- and ST-HSCs. MPPs number, instead, was unchanged and their frequency decreased accordingly (Fig. 1E and F). Consistently, the *in vivo* BrdU incorporation assay showed marked increase of proliferating NPMc+/YFP LSKs, LT-HSCs and ST-HSCs and, to a lesser extent, MPPs (Fig.S1A). No differences were observed in the frequency of apoptotic cells (Fig.S1B). The observed expansion of LT-HSCs in the pre-leukemic NPMc+ BM was unexpected, since previous studies using different cell-surface markers (e.g. SLAM(24)) and the knock-in *Npmc+* mouse model reported equal(25) or slightly decreased(6) HSCs frequency/numbers. Thus, we analyzed NPMc+ and control LSKs from our mouse model using the SLAM markers (see Fig.S1C for the FACS gating strategy), and found same numbers and a slightly decreased percentage of LSK/CD150+/CD48- HSCs, similar to previous studies (Fig.S1D, upper panels). However, analysis of the CD150+CD48- populations using the CD34 marker, which distinguishes HSCs with long term self-renewal (CD34^-^ HSCs) from early multipotent progenitors (CD34^+^-MPP1)(26), showed increased numbers and frequency of CD34^-^ HSCs (fig.S1D, lower panels), thus confirming the expansion of LT-HSCs induced by NPMc+.

We next investigated whether NPMc+ - dependent HSC proliferation correlates with expanded self-renewal *in vivo*, by measuring numbers of donor-derived LT-HSCs after transplantation of equal numbers of LT-HSCs (the experimental scheme is depicted in Fig. 1G). Unfortunately, *in vitro* culturing of BM-MNCs with cytokines, as required for TAT-CRE mediated expression of NPMc+, modulates expression of lineage markers on HSCs(27). Importantly, this modulation does not modify their repopulating ability (thus not affecting the data presented above)(27), yet it prevents the quantitation of donor LT-HSCs prior transplantation, as required to evaluate their self-renewal quotient. To circumvent this limitation, we crossed the NPMc+/YFP mice with the CMV-CreER^T^ strain(28) to generate NPMc+/YFP/CRE and YFP/CRE (as control) animals, where NPMc+ and YFP expression can be rapidly induced *in vivo* upon 7-day treatment with 4-OH-tamoxifen. YFP+ BM-MNCs were purified from 4-OH-tamoxifen-treated NPMc+/YFP/CRE or YFP/CRE mice and injected into lethally irradiated syngeneic mice (BM inputs). Though numbers of donor LT-HSCs were comparable in the NPMc+/YFP/CRE and YFP/CRE samples (144.8±17.62 *vs* 120.1±16.19 in 10^6 BM-MNCs, respectively; p=0.15; Fig. 1H), animals transplanted with NPMc+/YFP/CRE cells showed a significantly higher number of LT-HSCs at four months after transplantation (416.4±126.7 *vs* 190.6±82.4; p=0.0003; Fig. 1H) and, accordingly, a significantly higher self-renewal quotient(29) (45.2±2.84 vs 24.9±2.23; p=0.01; Fig. 1I). To investigate whether expanded NPMc+ HSCs preserve their repopulating abilities, we performed a limiting dilution transplantation. As shown in Fig. 1L, the NPMc+/YFP/CRE BM-MNCs (recovered four months after BMT) showed a significantly higher HSCs frequency evaluated as Competitive Repopulating Units (CRUs) by ELDA software analysis(30), as compared to YFP/CRE BM-MNCs (∼1:9,000 *vs* ∼1:20,000; p<0.05), thus confirming, functionally *in vivo*, that NPMc+ expression increases HSC self-renewal and that the expanded pool of NPMc+_HSCs is fully competent in repopulating the mouse bone marrow upon secondary transplantation.

### Increased proliferation by NPMc+ does not reduce the pool of quiescent HSCs

HSCs are largely quiescent, a phenotypic state that is critical for the maintenance of lifelong self-renewal. Since recruitment of HSCs into the cell cycle is associated with the reduction of quiescent HSCs and progressive HSCs exhaustion (31), (32), we evaluated the effect of NPMc+ on LT-HSCs quiescence, by combined analyses of DNA-content and expression of the Ki67 proliferation-associated antigen (Fig. 2A, left panel). Results confirmed the increased fraction of cycling NPMc+ LT-HSCs in the Ki67+ population (Ki67+/>2N; S/G2/M population Fig. 2A, middle panel). Surprisingly, however, we did not observe any decrease in the percentage of quiescent cells within the LT-HSC compartment (G0 cells, Fig. 2A, middle panel). Notably, since overall the number of LT-HSCs is expanded in the NPMc+ BM (as shown in Fig. 1F), also numbers of G0 HSCs were proportionally expanded in the total BM-MNCs (551.3+/-137.8 in one million of NPMc+/YFP BM-MNCs vs 169.7+/-52.6 in the control; p<0.02) (Fig. 2A, right panel). The expanded number of quiescent LT-HSCs in the NPMc+ BM was confirmed by the BrdU-based pulse-chasing Labeling Retaining Assay(26) (LRA) (Fig. 2B, upper panel), which revealed higher numbers of label-retaining BrdU+ LT-HSCs in the presence of NPMc+ expression (Fig. 2B; lower panel). Finally, to investigate whether quiescent NPMc+ LT-HSCs are fully competent in supporting hematopoiesis, we measured the rate of mouse survival after *in vivo* treatment with the cell cycle specific 5-fluorouracil (5-FU) drug. In fact, upon weekly injection of 5-FU, mouse survival depends on the capacity of quiescent LT-HSCs to re-enter the cell cycle and reconstitute hematopoiesis(33). Mice reconstituted with NPMc+/YFP were significantly more resistant to 5-FU treatment than control YFP mice (Fig. 2C), thus demonstrating that quiescent NPMc+ HSCs are functionally competent in supporting hematopoiesis.

**Figure 2.**
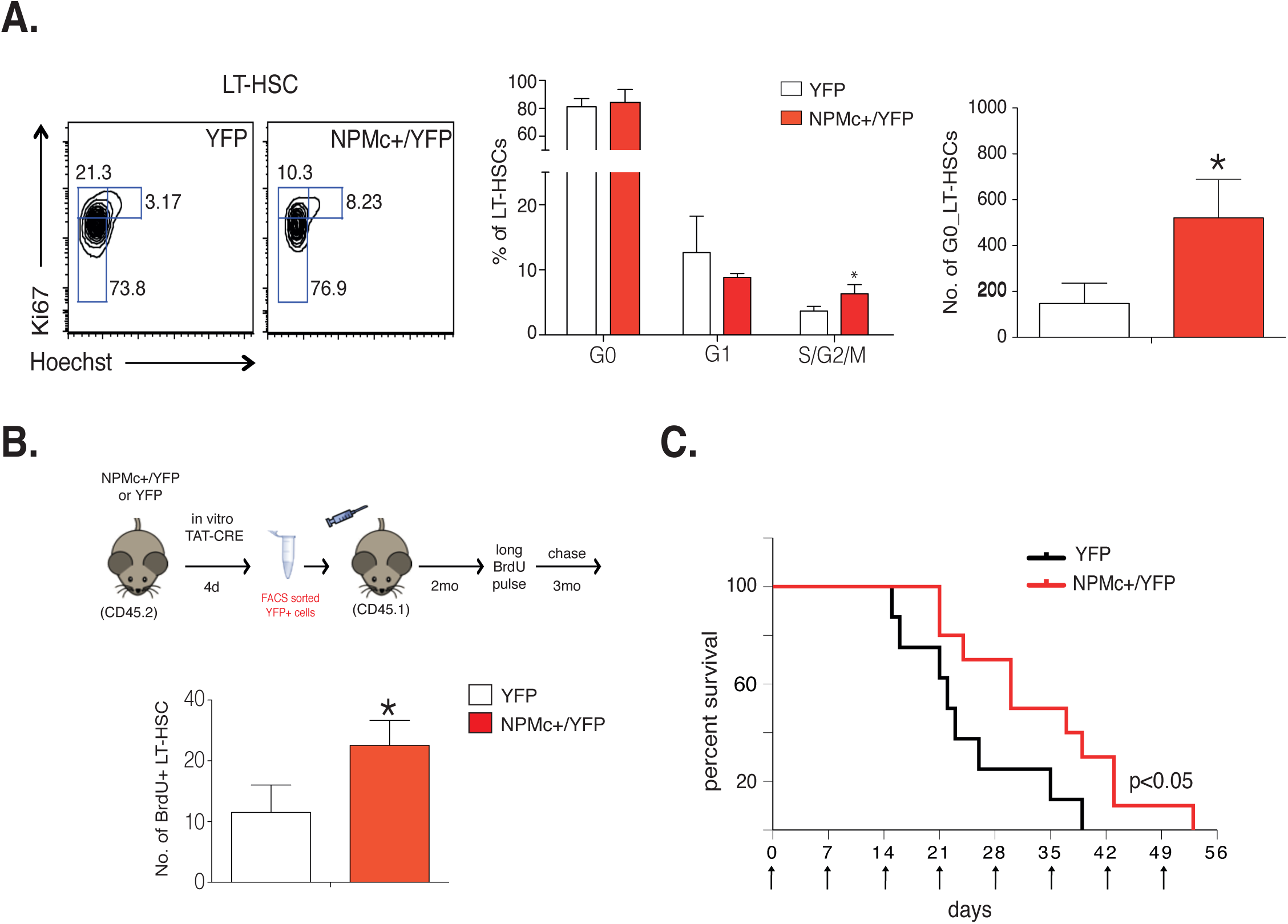
NPM1c+ expression preserves HSCs quiescence. **A.** Left panel: Representative plots showing the FACS gating strategy for the cell cycle analysis of LT-HSCs, based on Hoechst/Ki67 double staining. Middle panel: percentage of LT-HSCs in the different cell-cycle phases determined by FACS analyses (4 animal/group; *=p<0.05). Right panel: numbers (per million of BM-MNCs) of quiescent (G0) LT-HSCs in the same animal cohorts. **B.** Top: schematic representation of the experimental approach to quantitate BrdU+ label retaining cells (LRCs). Bottom: numbers of BrdU+ LR LT-HSCs per million of BM-MNCs detected at the end of the chase period. (4 animals/group, graph representative of 1 of 2 independent experiments; *=p<0.05). **C.** Kaplan Meyer survival curve of YFP and NPMc+/YFP mice weekly treated with 5-FU (8 animals per cohort, graph representative of 1 of 2 independent experiments). The reported p value was calculated with the log rank test.

Collectively, these data show that NPMc+ expression exerts a dual effect on LT-HSCs: on one side it induces proliferation and expansion, on the other, it maintains quiescence and preserves their self-renewal potential.

### NPMc+ restores quiescence and prevents Flt3-ITD HSCs exhaustion

It has been shown that *Flt3*-ITD LT-HSCs hyper-proliferate, are decreased in numbers and have impaired self-renewal(34), suggesting progressive exhaustion of the HSC pool due to constitutive proliferative/differentiative signals delivered by *Flt3*-ITD (32, 35, 36). Consistently, *Flt3*-ITD knocked-in mice do not develop AML(34), unless they are crossed with animals carrying a cooperative mutation, such as NPMc+ (3, 4). We thus investigated whether co-expression of NPMc+ rescues the defective *Flt3*-ITD HSC-phenotype. To generate NPMc+/*Flt3-ITD* compound mice, we first crossed our transgenic conditional NPMc+ mice with mice carrying the *Cre* recombinase under the control of the poly I:C (pIpC)-inducible Mx1 promoter (MxCre mice(37)), and then with a mouse strain carrying the *Flt3-ITD* knock-in mutation (*Flt3*^+/ITD^ mice(38)). The resulting NPMc+/MxCre (thereafter NPMc+/Mx) and NPMc+/*Flt3*^+/ITD^/MxCre (thereafter NPMc+/Flt3-ITD/Mx) were treated with pIpC to induce NPMc+ expression, while pIpC treated MxCre (thereafter Mx) and *Flt3*^+/ITD^/MxCre (thereafter Flt3-ITD/Mx) were used as controls.

After ten days of *in vivo* pIpC treatment, NPMc+ was expressed in most PB WBCs (Fig.S2 A-B) and LSK cells of both NPMc+/Mx and NPMc+/Flt3-ITD/Mx mice (Fig.S2C). According to previous reports(3, 4), NPMc+/Flt3-ITD/Mx mice developed fully penetrant AMLs with short latency (median of 69 days; Fig.S2D). Numbers of HSCs and early progenitors were measured by FACS analyses in BM-MNCs at three weeks after pIpC treatment. LT-HSCs were increased in the NPMc+/Mx mice (confirming data obtained with NPMc+/YFP+ mice in Fig. 1F) and significantly reduced in the Flt3-ITD/Mx, as reported(34) (Fig. 3A). Strikingly, expression of NPMc+ rescued the reduced number of LT-HSCs observed in the BM of Flt3-ITD/Mx mice (Fig. 3A). Moreover, NPMc+/Flt3-ITD/ Mx mice showed significant expansion of ST-HSCs, MPPs and LSKs numbers in the BM when compared to Flt3-ITD/Mx animals (Fig. 3A). Thus, the expression of NPMc+ rescues the depletion of HSCs in the Flt3-ITD/Mx bone marrow.

**Figure 3.**
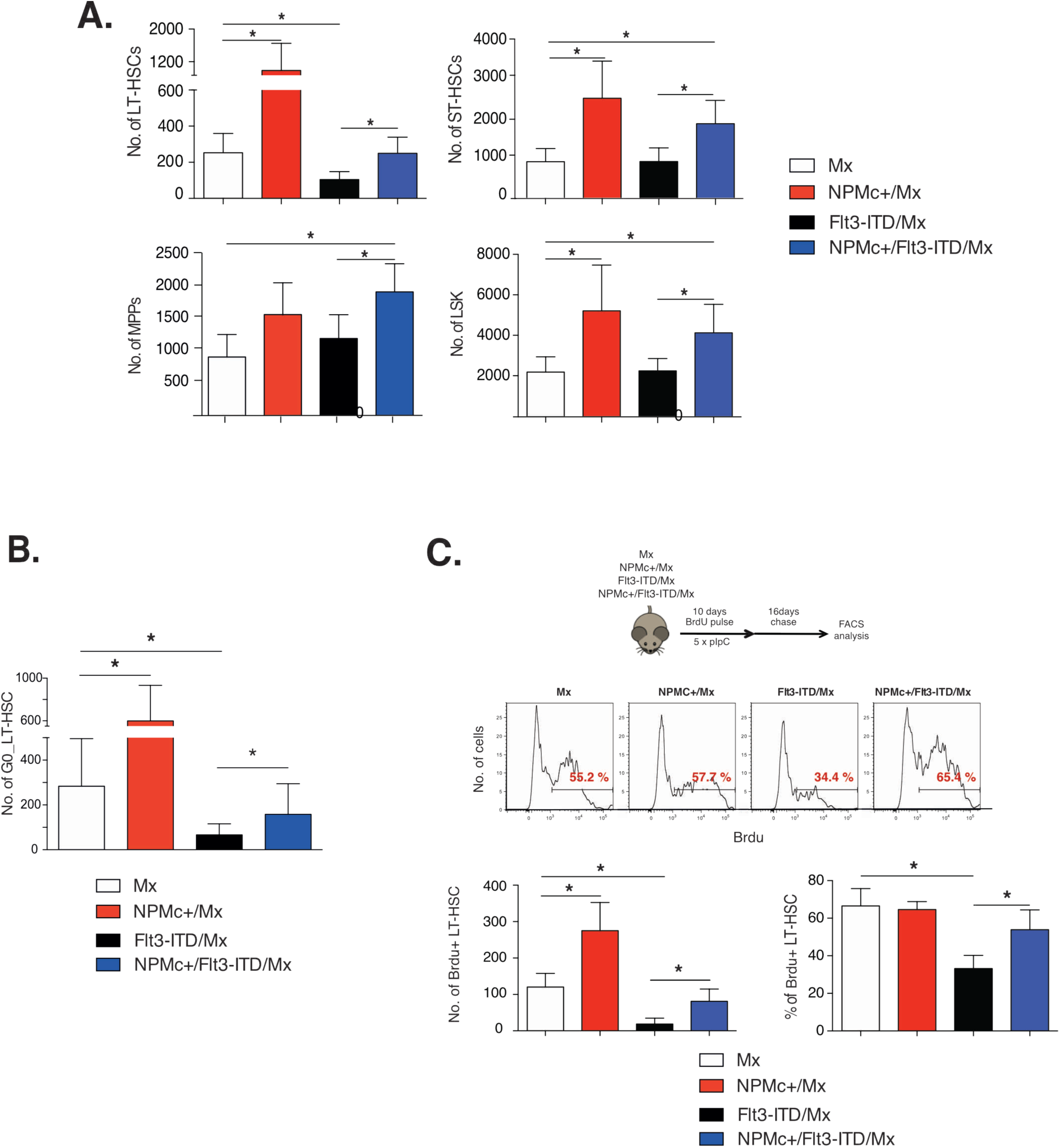
NPMc+ expression restores numbers of total and quiescent LT-HSCs in Flt3-ITD mice. **A.** Numbers of HSC/progenitors per million/BM-MNCs in Mx, NPMc+/Mx, Flt3-ITD/Mx and NPMc+/Flt3-ITD/Mx mice (pool of 3 independent experiments; *p<0.05). **B.** Cell cycle analysis by Hoechst/Ki67 staining: numbers, per million of BM-MNCs of quiescent (G0_2N/Ki67-) LT-HSCs (12 animal per cohort; pool of four independent experiments). **C.** Upper panel: Experimental scheme of the BrdU pulse-chasing retaining assay. Middle panels: gating strategy for the analyses of the BrdU retaining LT-HSCs after 16 days of chasing. Lower panels: numbers per million of BM-MNCs (left) and percentages within the HSC compartment (right) of BrdU+ LT-HSCs in NPMc+/Mx (n=5), Flt3- ITD/Mx (n=5), NPMc+/Flt3-ITD/Mx (n=6) and control Mx-CRE (n=5) mice (*=p<0.05).

We then investigated whether exhaustion of Flt3-ITD/Mx HSCs is associated with a reduction of quiescent HSCs and whether NPMc+ co-expression induces their re-expansion. HSC quiescence was measured by FACS analyses of LT-HSCs stained with the anti-Ki67 antibody. Numbers of G0 LT-HSC in the BM-MNCs, as observed for total LT-HSCs (Fig. 3A), were significantly expanded in the NPMc+/Mx mice, dramatically reduced in the Flt3-ITD/Mx mice and re-expanded in the NPMc+/Flt3- ITD/Mx, to an extent that was comparable to control Mx mice (158.5±135.9 vs 283.8±211.9/10^6 BM-MNCs, respectively; p=0.09. Fig. 3B lower panel). Then, to obtain an independent confirmation of the effects of NPMc+ and *Flt3*-ITD on HSC quiescence, we measured numbers and frequencies of label-retaining cells, using an LRA “shortened” protocol to fit the reduced pre-leukemic phase of NPMc+/Flt3-ITD/Mx mice (see Fig. 3C, upper panel for experimental design). FACS analyses of BM- MNCs showed a significantly decreased number of quiescent/slowly replicating BrdU+ LT-HSCs in the Flt3-ITD/Mx sample (Fig. 3C, bottom left panel), in agreement with the observed reduced numbers of G0 LT-HSCs (Fig. 3B, lower panel) and consistent with the reported ability of *Flt3*-ITD to increase HSCs proliferation(34). Notably, co-expression of NPMc+ restored numbers of BrdU+ LT- HSCs to the levels of control animals (Fig. 3C, bottom left panel). If we consider the percentage of BrdU+ cells within the LT-HSCs compartment, again we observed no variation in NPMc+/Mx mice (as in Fig. 3B, upper panel) and a significant decrease in Flt3-ITD/Mx mice that was rescued by NPMc+ co-expression (Fig. 3C, bottom right panel, and middle panels as representative results). These data demonstrate that NPMc+ co-expression preserves quiescence and numbers of Flt3-ITD HSCs, thus preventing their functional exhaustion.

### NPM1c+ expression enforces a quiescence transcriptional program

We then investigated whether the biological effects of NPMc+ on HSCs quiescence correlate with the activation of a specific transcriptional program. Gene-expression analyses of NPMc+/YFP+ *vs.* YFP+ LT-HSCs showed up-regulation of *HoxA* genes in this compartment (Fig. 4A), which are critical NPMc+ targets and a hallmark of NPMc+ AMLs(39, 40). Gene Set Enrichment Analysis (GSEA) of genes differentially expressed in the two samples (Table S1A) showed enrichment, in NPMc+ LT-HSCs, of transcriptional programs specific of HSCs (Fig.S3A-B), NPMc+ or rMLL AMLs (Fig. S3C-D and S3E) and LSCs (Fig.S3F). Most notably, we also found significant enrichment of genes up-regulated in quiescent HSCs(41) (Fig. 4B, left panel) and upregulation of genes that are essential for the maintenance of HSC self-renewal through enforcement of quiescence (e.g., *Mpl, Cdkn1a, Tgfb, Egr1 and Angpt1*)(42) (Fig. 4C). Thus, NPMc+ imposes a transcriptional program in wild-type LT-HSCs that is consistent with its ability to promote both quiescence and self-renewal.

**Figure 4.**
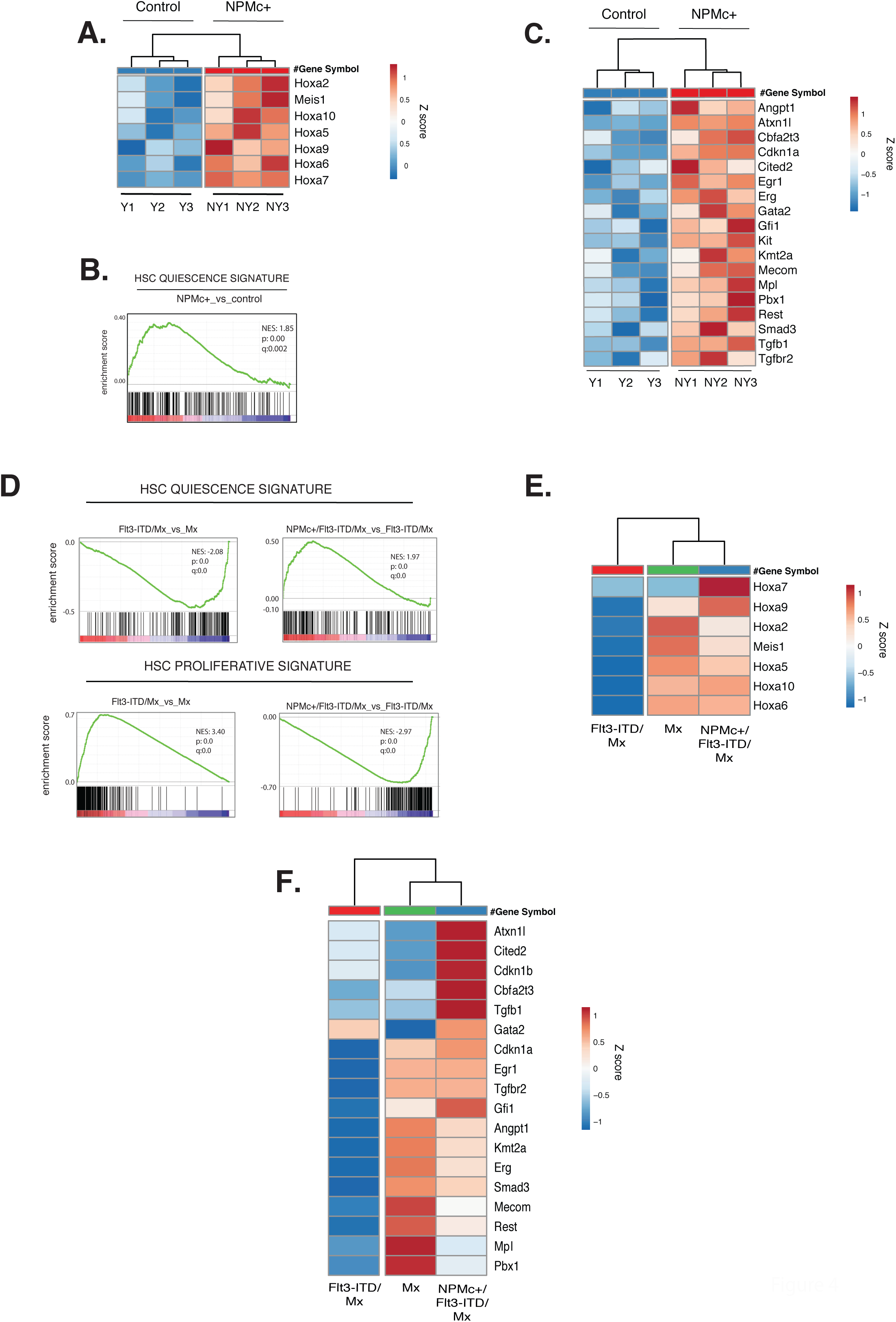
NPM1c+ expression enforces a quiescence transcriptional program. **A/C.** Gene- expression analyses of FACS sorted YFP (Y) and NPMc+/YFP (NY) LT-HSCs (4 animals per pool, 3 independent experiments). The heatmaps show the Z-scores of normalized expression values of each replica for homeobox genes (A) and genes involved in HSC quiescence maintenance (C); clustering using Euclidean distance and average linkage. **B/D**. GSEA for HSC quiescence- and proliferative-signatures in NPMc+ vs. control (B), Flt3-ITD/Mx vs Mx and NPMc+/Flt3-ITD/Mx vs Flt3- ITD/Mx (D) LT-HSCs. Normalized enrichment score (NES), p value (p) and false discovery rate (q) are indicated. **E/F**. As in A/C, for homeobox genes (E) and quiescence genes (F) (each column represents the average value of two independent experiments for Flt3-ITD/Mx; Mx and NPMc+/Flt3-ITD/Mx samples).

GSEA of genes differentially expressed in Flt3-ITD/Mx *vs.* Mx LT-HSCs control animals (Table S1B) showed depletion of quiescence genes and enrichment of proliferation genes in the Flt3-ITD/Mx HSCs, both rescued by NPMc+ co-expression (Fig. 4D). Similarly, HSCs/LSCs expression signatures were down-regulated by Flt3-ITD and rescued by NPMc+ (Fig.S3 G and H). At single-gene level, several of the self-renewal/quiescent genes up-regulated by NPMc+ in LT-HSCs were markedly downregulated by Flt3-ITD and rescued by NPMc+ (e.g. *Hoxa5, Hoxa10, Hoxa6, Cdkn1a, Egr1, Smad3, Tgfbr2*), while others, though not affected by Flt3-ITD, were up-regulated in the double-mutant LT-HSCs (e.g. *Hoxa7, Cited2, Cdkn1b, Tgfb1*) (Fig. 4E-F). In conclusion, NPMc+ enforces its transcriptional program upon Flt3-ITD/Mx LT-HSCs, restoring expression of critical HSC quiescence genes.

### NPMc+ supports both dormant and active states of quiescent HSCs, and induces their co-expression in the Flt3-ITD HSCs

We next investigated how the effects of NPMc+ on quiescence protects *Flt3*-ITD HSCs from functional exhaustion, and how they cooperate to induce rapid transformation of HSCs. Quiescent HSCs include two phenotypically distinct sub-populations: i) dormant HSCs (dHSC), which divide very unfrequently (approximately five divisions *per* life time), have the highest self-renewal potential, and are activated by stress signals; and ii) active HSCs (aHSC), which, though quiescent, are prone to divide and support homeostatic hemopoiesis (26). Notably, HSCs can switch dynamically between the two phenotypic states through a series of transient intermediate cell states(43, 44). Thus, we analysed whether expression of NPMc+ and/or *Flt3*-ITD alters “dormancy” versus “activity” equilibrium within the pool of quiescent HSCs.

Dormant and active HSCs can be transcriptionally defined by single-cell (sc) RNA-seq analyses, through their position within a pseudo-temporal trajectory(43). Pooled scRNAseq data from our four samples (10,519 cells) were ordered along a pseudo-temporal trajectory using the MolO dHSC gene-signature(43, 45) (Fig. 5A left panel). Consistently, the trajectory was characterized by the gradual decrease, from left to right, of the Normalized Mean Expression (NME) of the MolO signature (Fig. 5A, right panel). We further observed the decrease of a second, independent dHSC signature (Do28; Fig.S4A, TableS2A), and the concomitant gradual increase of signatures specific for aHSCs (Act166), G1-to-S progression or DNA-replication (Fig.S4 and TableS2A). Thus, our pseudo-temporal trajectory is able to identify the two HSCs states (dormant and active) and cells transiting between the two.

**Figure 5.**
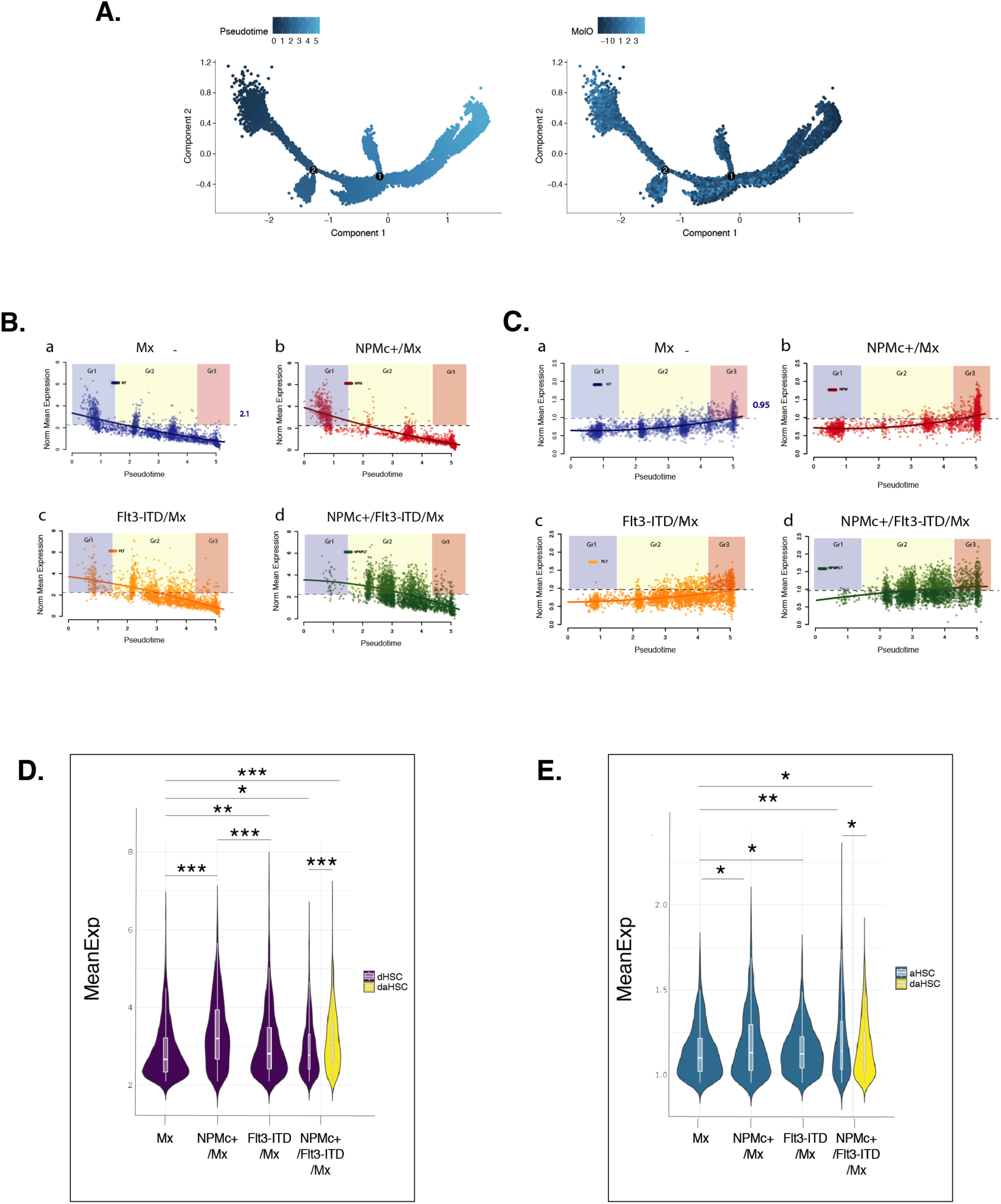
NPMc+ expression promotes HSC dormancy. **A.** Left panel: pseudo-time trajectory of pooled scRNA seq data (Mx, NPMc+/Mx, Flt3-ITD/Mx, NPMc+/Flt3-ITD/Mx) depicting the transition from the dormant to the active state of LT-HSCs; pseudo-time progression is represented by the color-code dark (dormant) to light blue (active). Right panel: color-coded normalized mean expression (NME) of the MoLO gene-signature along the pseudo-time trajectory. High MoLO expression (light blue) defined dormant state while low MoLO expression (dark blue) define the active state. **B-C**. scatter plots of NME values of dormant (B) and active (C) genes by pseudotime in the 4 samples (sig_Do and sig_Act gene-lists; TableS2B). Colored lines are polynomial fit of data and depict the trend of the NME of each signature in each sample. In panels B-a and C-a the horizontal dotted line defines the 99th percentile of the NME distribution of Mx cells in Group 3 and 1, respectively. Cells above the thresholds are dHSC (B) or aHSC(C) and are framed by semi-transparent colored squares, based on groups definition. **D-E**. Combination of violin- and overlaid box-plots represent the NME distributions of: sig_Do genes (D) or sig_Act genes (E) in the dHSCs or aHSCs respectively of all samples (only for the NPMc+/Flt3-ITD/Mx sample dHSCs, aHSCs and daHSCs are depicted separately). Statistical significance was tested with one tailed Wilcoxon rank sum test. *p<0.05; **p<10-4; *** p<10-10

We then set the experimental conditions allowing for the identification of dHSCs and aHSCs in the trajectory, and the extent of activation of dormant and active transcriptional programs for each cell. Briefly (see Methods for details), cells of each pseudo-temporal trajectory were divided in three consecutive groups (Groups1-3 from left to right; Fig.S4B), with Group1 and 3 enriched for dHSC and aHSC, respectively, and Group2 comprising cells transiting between the two states. We have identified dormant and active genes that are differentially expressed among Groups and samples (TableS2B), and plotted their NME values along the pseudo-time of each sample (Fig. 5 B and C for dormant and active genes, respectively). We then set the dormant and active NME cut-off values above which we can define dHSC and aHSC (Fig. 5B and C panels a, respectively), with cells below both cut-off values being considered as transiting among the two phenotypic states (trHSCs). We applied the same thresholds to all four samples (fig. 5 B and C panels b,c and d) and, for each of them, we calculated numbers and frequencies of dHSCs, aHSCs and trHSCs (as detailed in TableS2C) and the mean average expression of dormant and active gene signatures (Fig. 5 D and E).

NPMc+/Mx LT-HSCs showed increased numbers of aHSCs (35,4% vs 15,1% in control cells) and a shift of the pseudo-time trajectory toward its right-end, as evidenced by the expansion of Group3-aHSCs (30% vs 9.3% in control cells) and the reduction of Group2-trHSCs (18,4% vs 39,3% in control cells) (Fig. 5B,C; TableS2C). This accelerated transition toward aHSCs correlates with increased expression of active genes (Fig. 5E), yet was not accompanied by the depletion of Group1-dHSCs, which showed instead a slight expansion (20,2% vs 18,4% in control cells, TableS2C), and correlates with a significantly higher expression of dormant genes (Fig. 5D). In summary, NPMc+ exerts the dual function of favouring HSCs transition toward the active state, eventually leading to increased proliferation (as shown in fig. 2A, middle panel) and preserving an intact pool of dormant HSCs, thus explaining, mechanistically, its capacity to support high self-renewal potential *in vivo*. Noteworthy, this experimental setting does not allow to appreciate the expanded number of NPMc+_HSCs in the bone marrow (previously defined by FACS analysis of the whole bone marrow population), since scRNAseq were performed on same number of purified LT-HSCs cells to evaluate variations of HSC sub-populations.

The trajectory of Flt3-ITD/Mx HSCs was characterized by a marked mobilization of dHSCs from Group1 (7,3% vs 18,4% in control cells) to Group2 (24,2% vs 11,5% in control cells) in the absence of gross variations in the total number of dHSCs (32,4% vs 30,1% in control cells). aHSC were moderately expanded (19.6% versus 15% in control), slightly shifted toward Group3-aHSCs (13% versus 9.6% in control) and showed, accordingly, a mild upregulation of active genes (Fig. 5E). Thus, Flt3-ITD induces irreversible commitment of dHSCs toward a more active state which can result into reduced self-renewal and progressive exhaustion, as observed *in vivo*.

The NPMc+/Flt3-ITD/Mx pseudo-time trajectory showed pronounced accumulation of HSCs in Group2 (80.1% versus 56% in control cells, TableS2C), accompanied by the almost complete depletion of Group1-dHSC (1.1% vs 18.4% in control cells), and increased numbers of both aHSCs (49,4% vs 15,1% in control cells) and dHSCs (58,3% vs 30,1% in control cells). Virtually all dHSCs (more than 90%) and most aHSC (about 80%) were localized in Group2, suggesting an overlap of the two phenotypic states at single-cell level. Indeed, while in Mx, NPMc+/Mx and Flt3-ITD/Mx samples the dormant and active states are mutually exclusive, NPMc+/Flt3-ITD/Mx HSCs showed a high percentage of cells displaying, at single-cell level, both dormant and active states (daHSCs; 32% vs 0,4% in control cells, mainly located in Group2) (TableS2C). The HSCs with this “new” phenotypic state showed increased expression of both dormant and active genes, as compared to control LT-HSCs (Fig. 5D,E) and, remarkably, higher expression of dormant genes also when compared to dHSCs in the same sample (fig. 5D). These data demonstrate that the NPMc+ and Flt3-ITD cooperation is the result of the integration of their pathogenic signals within a new HSCs phenotypic state that, although forced by Flt3-ITD to exit the “physiological” Group1 dormant-state and progress toward the active state, is still driven by NPMc+ to express a “dormant-like” status embedded in the same active-committed cell. This unique state generates a pre-leukemic HSC that can, in principle, proliferate without loosing its self-renewal potential.

### Pharmacological inhibition of the dormancy-related TGFβ pathway in NPMc+/*Flt3-ITD* AML reduces self-renewal of LSCs

Finally, we investigated the functional role of dormancy in the regulation of self-renewal in leukemia SCs (LSCs). TGFβ1 promotes cellular dormancy(46, 47), and is one of the actionable genes upregulated by NPMc+ in both control and /Mx LT-HSCs. Consistently, TGFβ signaling is activated in these cells (Fig. 6A-B). To investigate the effect of TGFβ1 inhibition on self-renewal of LSCs, we analyzed the repopulating potential of leukemic cells after *in vivo* treatment with the LY364947 TGFβR-I inhibitor (see Fig. 6C for the experimental design). Levels of engraftment at 10-15 days post transplantation were comparable between the animals transplanted with blasts pre-treated or not with the inhibitor. However, starting from 30 days post transplantation, mice transplanted with LY364947-treated blasts showed progressive decline of blasts in the PB (Fig. 6D) leading to mouse survival (p<0.01; Fig. 6E and Fig.S5 B and C for an independent *NPMc+/Flt3-ITD* AML experiment). LY364947 treatment of mice with growing *NPMc+/Flt3ITD* leukemia significantly prolonged their survival (Fig.S5A). Thus, the NPMc+ -dependent and quiescence-related TGFβ pathway identified in the pre-leukemic HSCs is critical for LSCs self-renewal (Fig. 6D), and pharmacological inhibition of this pathway slows down disease progression (Fig.S5A), resulting into significantly increased mouse survival.

**Figure 6.**
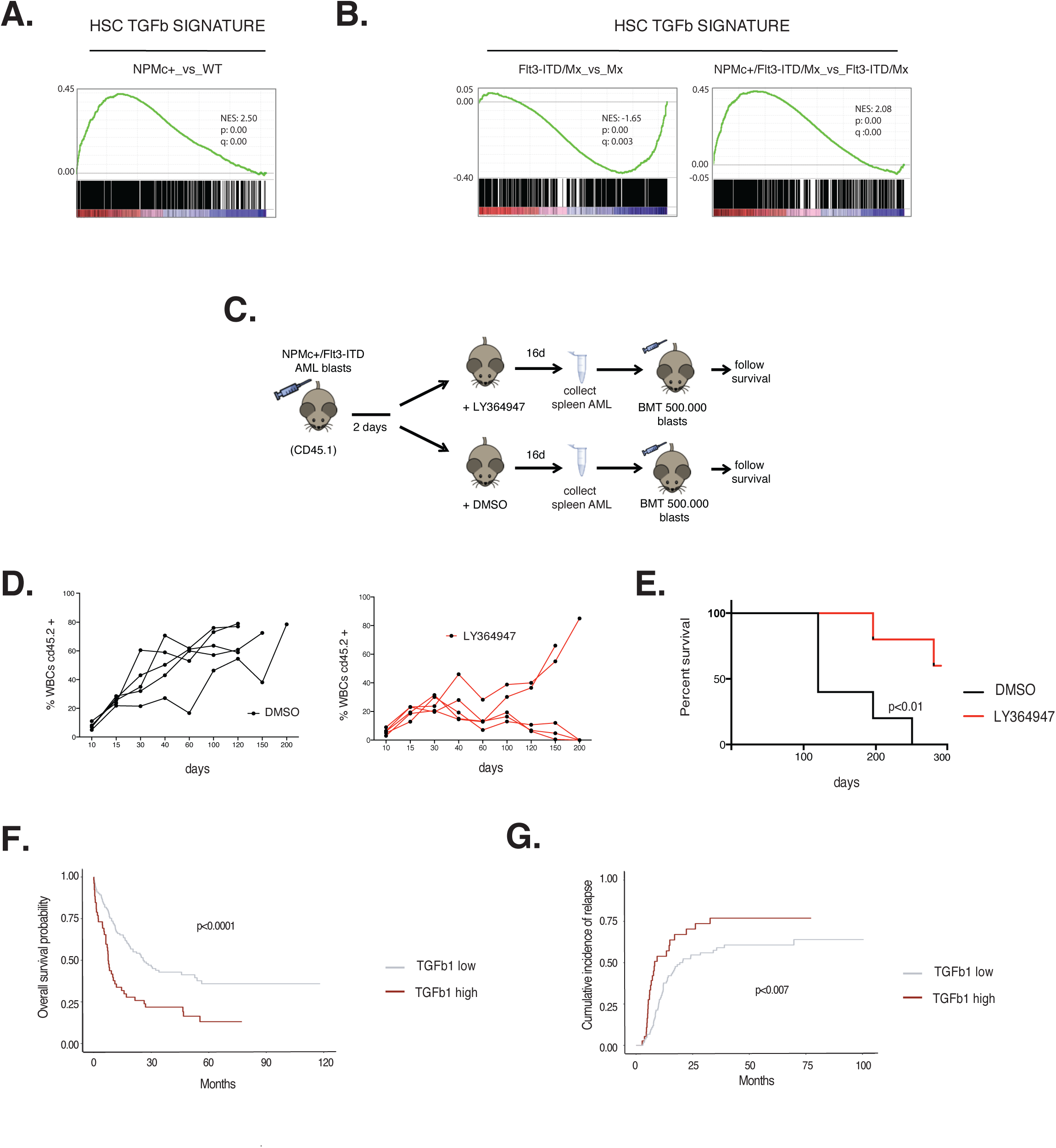
Pharmacological inhibition of the TGFβ pathway affects *NPMc+/Flt3-ITD* LSC maintenance. **A-B.** GSEA enrichment plots for the TGFβ signature (70) in NPMc+/YFP LT-HSCs vs YFP LT-HSCs (A) or Flt3-ITD/Mx vs Mx LT-HSC and NPMc+/Flt3-ITD/Mx vs Flt3-ITD/Mx LT-HSCs (B). Normalized enrichment score (NES), p value (p; log rank test) and false discovery rate (q) are indicated. **C.** Experimental scheme: mice transplanted with a *NPMc+/Flt3-ITD* AML blasts were treated with LY364947 or vehicle every other day for 16 days. At the end of the treatment 500,000 blasts were purified and re-transplanted in secondary recipient mice. **D.** Percentage of control or LY364947 treated blasts (CD45.2+) in the PB of transplanted (CD45.1+) mice (5 animals per group) **E.** Kaplan Meier survival curve of mice transplanted with LY364947-treated or control blasts (5 animals per group). **F.** Overall survival and cumulative incidence of relapse (**G**) of TCGA AML patients based on the expression of *TGFβ1* (normalized RSEM, high>=0.532, low<0.532).

Finally, we preliminarily investigated the clinical value of this observation by assessing the impact of *TGF*β*1* expression on overall survival (OS) and cumulative incidence of relapse (CIR) of AML patients, using the TCGA cohort of 200 AML-cases(48). Best *TGF*β*1* expression cutoff-predictor of OS was identified by ROC analyses using clinical and RNAseq data from 173 AML samples (Fig.S5D). Patients with high *TGF*β*1* expression (n=52) showed significantly shorter OS (p<0.0001) and higher CIR (p<0.007) than patients with low *TGF*β*1* expression (N=121) (Fig. 6F and G). Most notably, Cox multiple-regression analyses showed that high *TGF*β*1* expression is an independent predictor of OS and CIR, as compared to well-established prognostic factors(49) (including age, white blood cell counts and risk classes) (HR 1.88, 95% CI 1.27-2.80, p=0.002) (TableS3A-B). GSEA of genes differentially expressed between the two AML groups (TGFβ1-high versus TGFβ1-low, Table S4) showed enrichment, in the TGFβ1-high samples, of TGFβ (Fig.S5E), *NPMc+* leukemia (Fig.S5F) and dormancy (MolO and Do28; Fig.S5G) transcriptional programs; conversely, no enrichment was found for the aHSC gene signature Act166 (Fig.S5G). Analyses of the distribution of cases with high or low *TGF*β*1* expression across the different cytogenetic and molecular types of AMLs showed a trend toward significant association between high *TGF*β*1* expression and the presence of *NPM1* mutations (TableS3C), however, cases with high *TGF*β*1* expression were also found among other molecular types, including AMLs with complex karyotypes, recurrent translocations or *FLT3* mutations, suggesting that increased TGFβ signaling and enforcement of the dormancy program is a general feature of AMLs with poor prognosis.

## Discussion

We investigated cell cycle and self-renewal properties of pre-leukemic and leukemic SCs expressing NPMc+ and/or *Flt3-ITD*, the most frequent co-occurring mutations in AML patients(8). First, we have shown that NPMc+ expression has transcriptional and functional effects on the HSCs compartment. NPMc+ - expressing HSCs were increased in numbers, displayed higher BM repopulating potential and expanded self-renewal *in vivo*, strongly supporting the ability of NPMc+ to confer a pre-leukemic phenotype upon HSCs. Emerging evidence suggest that *NPMc+* may provide oncogenic functions in different hematopoietic compartments during AML clonal development also based on the occurrence of cooperative mutations. In AMLs with CHIP, mutations of *NPM1* are frequently associated with *DNMT3A* mutations and found in myeloid progenitors, where they induce extended self-renewal and HSC-reprogramming, thus favoring leukemia progression(10). In *de novo* AMLs, instead, mutations of *NPM1* are more frequently associated with *FLT3-ITD* and a stem-like phenotype (15), and can be found in the residual pre-leukemic HSC(18) . Moreover, *NPMc+* was found in 5-10% of myelodysplastic syndromes (MDS) and in the corresponding secondary AMLs(50), usually in the absence of *DNMT3a* mutations. Thus, *NPMc+* might be critical to establish a pre-leukemic phenotype by imposing aberrant self-renewal to either myeloid progenitors or HSCs, depending on the trajectory of AML development and associated co-mutations (CHIP or *de novo* AMLs, respectively). Recognition of *NPMc+* specific contribution in AML development may have important implications for patient prognostication and treatment. Indeed, the potential to target either HSCs or progenitors and the clinical implications have been previously demonstrated for the r-*MLL* AMLs, combining clinical and mouse model data(51–53). It is worth noting, that r-MLL and NPMc+ AMLs are molecularly similar, they depend on *HOX* genes expression to maintain self-renewal of Leukemic SCs (40, 54), can be treated pharmacologically by destabilizing the MLL complex(11, 55), and are characterized by transcriptional heterogeneity reflecting the cell-of-origin of the leukemia and allowing prediction of clinical aggressiveness and response to therapy(15, 19, 51).

We then showed that the effect of NPMc+ on self-renewal of HSCs involves reinforcement of a quiescence program. NPMc+ favors the expression of *HoxA* genes (as already shown in other compartments(3, 25)), a quiescent-HSC transcriptional program(41) and a number of genes known to preserve HSC self-renewal through promotion of quiescence (42). *Flt3-ITD*, instead, activates proliferation genes and down-regulates *HoxA* and quiescence genes, leading to depletion of quiescent HSCs and self-renewal impairment. NPMc+ enforces its transcriptional program upon *Flt3-ITD* HSCs, restoring expression of *HoxA* genes and HSC-quiescence genes and, accordingly, restoring the pool of quiescent HSCs and rescuing the defective phenotype of *Flt3-ITD* hyper-proliferating HSCs. These data support a model where oncogene-induced chronic hyper-proliferation of HSCs impairs self-renewal unless signals that increase quiescence (provided by NPMc+ in our model) are concomitantly present, thus allowing these mutations to be selected and to express their oncogenic potential.

We further investigated how the oncogenic signals delivered by NPMc+ and *Flt3-ITD* integrate and cooperate in HSCs, indeed, in our model, purified pre-leukemic NPMc+/*Flt3-ITD*/Mx HSCs induce AMLs upon transplantation (our unpublished data), suggesting that HSCs can be the cell of origin of NPMc+/*Flt3-ITD* AMLs. In particular, we analysed how NPMc+ and *Flt3-ITD* impact the dormant versus active state of quiescent HSCs. Indeed, quiescent HSCs can transit from a dormant to a more active state through a series of intermediate steps that describe a transcriptionally-defined pseudo-temporal trajectory and, to our knowledge, no studies have been reported aimed at investigating whether and how gene mutations involved in myeloid malignancies hijack this equilibrium. We showed that NPMc+ has a dual effect on quiescent LT-HSCs, enforcing both their dormant and active phenotypic-states. On one side, NPMc+ stimulates the transitions from dormant to active and eventually proliferative HSCs (as shown with *in vivo* cell cycle analysis); on the other side, enforcing a transcriptional program of dormancy, it stimulates re-entering of active HSCs into the dormant state, thus replenishing the pool of dHSCs and preventing exhaustion of their self-renewal potential. *Flt3*-ITD, instead, has milder effects on the dormant and active transcriptional states, but it induces an irreversible commitment of dHSCs toward a more active state, that, in the absence of a “re-entry” signal, leads to dHSC depletion and loss of self-renewal potential. In the presence of NPMc+, instead, the effect exerted by *Flt3*-ITD on HSCs leads to the full expression of its oncogenic potential. The constitutive signal to initiate the dormant-to-active transition elicited by Flt3-ITD, in the presence of enforced dormant and active states induced by NPMc+, creates a new phenotypic status where dormant and active states co-exist at single-cell level (dormant/active HSCs; daHSCs). We propose that in the dormant/active NPMc+/Flt3-ITD HSC, the acquisition of self-renewal or proliferation potential are not alternative fates, as it happens under physiological conditions, but co-exist within the same cell, thus allowing HSCs to proliferate indefinitely and to accumulate further genetic alterations, a condition that is permissive for the rapid selection of the leukemia-initiating SC. As recently shown, this strong NPMc+/*Flt3-ITD* cooperation at transcriptional level is based on a massive genome wide epigenetic rewiring(20).

Finally, we showed that pharmacological inhibition of dormancy decreases significantly self-renewal of LSCs in the context of the fully-expressed leukemic phenotype. Specifically, pharmacological targeting of TGFβ1, one of the actionable genes upregulated by NPMc+ and involved in the establishment of dormancy, reduced significantly the repopulating activity of transplanted NPMc+/*Flt3-ITD* murine AMLs and prolonged mouse survival, suggesting that dormant-related signals are integral to LSC extended self-renewal and leukemia outgrowth. Emerging evidence suggest that current treatment regimens, which are primarily designed to eliminate hyper-proliferating cells, might spare dormant pre-leukemic HSCs or LSCs, which are ultimately responsible for disease progression or relapse, respectively (56–58). Thus, the possibility to specifically target dormant related pathways may prevent the emergence of drug-resistant cells during primary treatment of *NPMc+* MDS or AMLs, or the expansion of minimal residual disease after therapy. Notably, we showed that high *TGF*β*1*-expression is an independent predictor of overall survival and incidence of relapse in AML patients across different molecular sub-types, including cases not expressing *NPMc+*, suggesting that enforcement of HSC dormancy might be shared by a number of other AML-associated mutations. Indeed, Li and colleagues reported that the AML-associated mutation *NRASG12D* enforces quiescence in a specific subset of HSCs, while it increases proliferation of a second HSC subset(59). In conclusion, our data suggest that increased *TGF*β*1* expression and enforcement of dormancy is a general feature of AMLs with poor prognosis, and that targeting TGFβ1/dormancy may represent a novel anti-leukemia therapeutic strategy.

## Materials and Methods

### Mice and treatments

The conditional *NPM1c+*(*3*) (hereafter NPMc+) and conditional Rosa26-*EYGF*(22) (hereafter YFP) strains were crossed to obtain NPM1c+/YFP and control YFP mice. Induction of YFP and/or NPMc+ expression *in vitro*: BM-MNCs derived from these mice were treated *in vitro* with recombinant TAT-CRE as reported(3). BM-MNCs/YFP+ were FACS sorted and injected i.v. into CD45.1 lethally irradiated recipient mice (8 Gy), and mice BM was analyzed 4 month after BMT.

In order to induce NPMc+ and/or YFP expression *in vivo*, NPM1c+/YFP and control YFP mice were crossed with a mouse strain expressing the *Cre-ER* transgene under the control of the CMV promoter (CMV-CreER^T^ (28), hereafter CRE) to obtain NPM1c+/YFP/CRE or YFP/CRE control mice. Both mice strains were injected i.p. with 1mg 4-OH-tamoxifen (Sigma) daily for 7 days to induce NPMc+ and/or YFP expression, then, BM-MNCs were purified and YFP+ cells were FACS sorted and used to perform the self-renewal assay (as described below).

In order to generate NPMc+/Flt3-ITD compound animals, we crossed our transgenic conditional NPMc+ mouse with a mouse strain carrying the *Cre* recombinase under the control of the Mx1 promoter (MxCre mice(37), hereafter Mx) and a mouse strain carrying the *Flt3-ITD* knocked-in mutation (*Flt3*^+/ITD^ mice(38), hereafter Flt3-ITD), to generate Mx, NPMc+/Mx, Flt3-ITD/Mx and NPMc+/ Flt3-ITD/Mx mice. To induce NPMc+ expression, animals have been injected every other day for ten days with poly(I)-poly(C) (250ug) (GE Healthcare). All the analyses were performed 21 days post-pIpC treatment.

Mice transplanted with AML blasts have been administered with TGFβR-I inhibitor LY_364947 (Selleckem) or vehicle alone diluted in 1xPBS at 10 mg/kg (of body weight) every other day.

5-FU was administrated at dosage of 150mg/kg by i.p., every 7 days.

Eight- to twelve-week-old mice were used throughout the study. All animal procedures were performed in accordance with the Italian Legislation. All mice were euthanized by high concentrations of CO2 inhalation.

### BrdU uptake and BrdU Label-Retaining assay

For the proliferation assay, 5-Bromodeoxy-uridine (BrdU, BD Bioscience) was administrated *in vivo* by i.p. injection (1mg/mouse) at two repeated doses every 6 h and mice were analyzed 12 h from the first injection. The BrdU Label-Retaining Cell assay (LRA) was performed as previously described(26).

### Purification of HSCs and Flow Cytometer analyses

BM-MNCs were isolated from mouse bones and stained for FACS analysis of membrane markers with the following antibodies (purchased from Bioscence): CD11b (PE-CY7) M1/70 clone; Ly-6G (PE- CY7) RB6-8C5 clone, Ter-119 (PE-CY7) Ter-119 clone, CD3e (PE-CY7) 145-2C11 clone, CD45R (PE-Cy7) RA3-6B2 clone, Ly-6A/E (PerCP-CY5.5) D7 clone, cKit (APC-eFluor780) 2B8 clone, FLK (PE) A2F10 clone, CD34 (FITC/biotinilated) RAM34 clone, streptavidin (eFluor450), CD45.1 (PE/FITC/APC) A20 clone, CD45.2 (PE/FITC/APC) 104 clone. Gating strategies are described in the results section.

For intracellular immune-staining (BrdU and Ki67), BM-MNCs were fixed in Cytofix/Cytoperm^TM^ buffer for 20min at room temperature (RT), washed by 1X Perm/Wash^TM^ Buffer (P/W) (BD Perm/WashTM buffer 10X diluted in dH2O) and re-fixed in Cytofix/Cytoperm^TM^Plus buffer for 10min at RT, protected from light. Cells were washed with P/W and incubated 5min with Cytofix/Cytoperm^TM^ buffer at RT, light protected, and washed again with P/W. Fixed cells were stained with anti-ki67 (Alexafluor647conjugated, clone 16A8, Biolegend) for 1hour at RT. After a final washing step the cells were stained with the DNA binding dye Hoechst 33342 (Sigma-Aldrich) at 4°C until FACS analysis. For BrDU staining, fixed and permeabilized cells (as described above) were treated with DNAse (150μg/ml) and then incubated with an anti-BrdU antibody (APC BrdU Flow kit, BD; dilution 1:100 in 1X Perm/Wash buffer) for 1 hour in ice.

### Transplantation assays

In the competitive setting, 1×10^6^ of CD45.2+ testing cells (YFP vs NPMc+/YFP) were mixed with 1×10^6^ competitive wild-type CD45.1+ BM-MNCs and injected i.v. into CD45.1+ lethally irradiated mice. At different time points up to 4 months after BMT, we evaluated the percentage of CD45.1+ and CD45.2+ in the PB-MNCs of mice.

In limiting dilution experiment, different amount of YFP or NPMc+/YFP sorted cells were mixed with 500,000 CD45.1 BM helper cells and injected i.v. in lethally irradiated recipient. Engrafted mice showed a frequency of YFP+ cells ≥0.1% in the peripheral blood at 4 months post transplantation. The frequency of LT-HSC has been calculated according to the ELDA software (http://bioinf.wehi.edu.au/software/elda).

### Immunofluorescence

For BM-MNCs immunofluorescence, cells were fixed in 4% PFA and spun onto microscope slide with the help of the Cytospin centrifuge. Staining with the anti-NPMc+ primary antibody (homemade rabbit polyclonal immunopurified ab, dilution 1:1000) was performed in a humid chamber for 4-5 hour at RT, followed by secondary antibody (Alexa488 Fluor® or Alexa647 Fluor®) and counterstained with DAPI. Samples were acquired with Leica TCS SP2 AOBS laser confocal scanner mounted on a Leica DM IRE2 inverted microscope (HCX PL APO 63X/1.4NA oil immersion objective) and analyzed by ImageJ 1.x.

For LKS immunofluorescence, cells were FACS sorted, and plated on poly-lysine treated coverslip. Cells were allowed to deposit by gravity, then were fixed in 4% PFA and stained as reported for BM-MNCs. For quantification of the NPMc+ signal, widefield images have been collected by Olympus BX61 fully motorized fluorescence microscope controlled by the image screening Scan^R software. The analysis has been carried out by a computational platform developed by our group (A.M.I.CO.)(60)

### RNA extraction and population-based RNA-seq

RNA was purified using QIAGEN RNeasy kit l from NPMc+/YFP and YFP FACS sorted LT-HSC and subjected to microarray analysis. Double stranded cDNA synthesis and related cRNA was performed with Nugen® Pico WTA Systems V2 (NuGEN Technologies, Inc). Hybridization was performed using the Affymetrix GeneChip® Mouse Gene ST 2.0 Arrays. Microarray data were normalized by RMA using Partek Genomic Suite 6.6.

For whole transcriptome sequencing, RNA was purified from FACS-sorted LT-HSCs with PicoPureTM RNA Isolation Kit (ThermoFisher). Sequencing libraries were generated using the SMARTseq protocol based on polyA-enrichment. 50bp single-end sequencing was performed with a HiSeq2000 device (Illumina). Sequences were aligned to the mouse reference genome (NCBI37/mm9) using TopHat2(61). After alignment, raw gene expression values were obtained with HTSeq(62). Differential gene expression was then estimated using the edgeR R package, using a TMM normalization(63).

### Single-cell library preparation and sequencing

For scRNA-seq libraries, we sorted LT-HSCs from pool of mice for each genotype. The libraries were prepared using the Chromium Single Cell 3ʹReagent Kits (v2): Single Cell 3ʹLibrary & Gel Bead Kit v2 (PN-120237), Single Cell 3ʹChip Kit v2 (PN-120236) and i7 Multiplex Kit (PN-120262) (10x Genomics) and following the Single Cell 3ʹReagent Kits (v2) User Guide (manual part no. CG00052 Rev C). Libraries were sequenced on NovaSeq 6000 Sequencing System (Illumina) with an asymmetric paired-end strategy (28 and 91 bp read length for R1 and R2 mate respectively) with a coverage of about 75,000 reads/cell.

### ScRNA-seq data analysis

Sequencing results were demultiplexed and converted to FASTQ format using Illumina bcl2fastq software. Sample demultiplexing, barcode processing and single-cell 3’ gene counting were obtained using a custom pipeline. The cDNA insert was aligned to the mm10/GRCm38 reference genome and associated with a gene using the GENCODE gtf file (downloaded from ftp.ebi.ac.uk//pub/databases/gencode/Gencode_mouse/release_M16/). Only confidently mapped, non-PCR duplicates with valid barcodes and unique molecular identifiers were used to generate the gene-by-cell matrix. Further analysis—including quality filtering, the identification of highly variable genes, dimensionality reduction, standard unsupervised clustering algorithms and a first sample and cluster markers identification— was performed using the Seurat R package analyzing both for single and merged samples.

After different quality filtering, we obtained 10,519 cells in total (2,481 for Mx, 2,332 for NPMc+/Mx, 2,385 for FLT3-ITD/Mx and 3,321 for FLT3-ITD/NPMc+/Mx). The mean and median numbers of detected genes-per-cell were 2,859.31 and 2,992.00 in WT, 3,156.01 and 3,190.00 in NPM, 2,880.96 and 2,980.00 in FLT and 2,879.66 and 2,901.00 in NPMFLT samples, respectively. We then normalized the raw data by the total expression of each cell and multiplied by a scale factor of 10,000.

To infer a different phenotypic state within our sorted LT-HSC cells and order them in a pseudotemporal trajectory we used Monocle 2 R package(64). The MolO list signature (n=27) was used as “ordering gene set” in this process. To obtain the trajectory we applied the “DDRTree” method for dimensional reduction with default parameters.

The pseudotime defined by the trajectory has been divided in three groups fixing two arbitrary thresholds considering the overall density profile of cells and the relationships between samples (e.g. values corresponding to the two main minima values of density in which samples intersect) (Fig.S4).

### Single cell RNAseq differential expression analysis

This “aggregated approach” (3 Groups of cells) allows for the characterization of the transcriptional profile not only at the extremities of the dormant-to-active transition process, but also at the intermediate state. The analysis tested each gene for differential expression by a multifactorial approach using as covariates *pseudotimes* – in terms of differences of expression between cells belonging to the different pseudotime groups, Gr1, Gr2 and Gr3 - and *samples*. In particular, we separately studied, pairwise, the following contrasts: NPMc+ vs Mx control, Flt3-ITD vs Mx control, NPMc+/Flt3-ITD vs Mx control and NPMc+/Flt3-ITD vs Flt3-ITD. In turn, each of these comparisons were challenged for differences in Gr2 vs Gr1, Gr3 vs Gr2 and Gr3 vs Gr1 groups of cells. These analyses were done with the differentialGeneTest Monocle’s main differential analysis routine that accepts a CellDataSet and two model formulae as input, and specifies generalized lineage models as implemented by the VGAM package. The final list of significant regulated genes (Sig_Do: dormant specific genes (Do28+MolO) differentially regulated in at least one of the contrasts. Sig_Act: active specific genes (Act166) differentially regulated in at least one of the contrasts -Table S2B-) was restricted to those that showed a raw count greater than 2 in at least 50 cells of the entire filtered datasets (10,519 cells) and that was regulated in at least one of the comparisons described above, with a *qval* value smaller than or equal to 0.01 (table S2B).

The Normalized Mean Expression for Sig_Do and Sig_Act genes was calculated for each cell along the pseudotime trajectory and used to define the criteria for the identification of dHSCs and aHSCs. In particular, we consider that in the pseudotime of normal Mx_HSCs most Group1-cells are dormant and, *vice versa*, most Group3-cells are active, therefore, we defined the threshold for cells expressing the dormant phenotype (dHSCs) as the 99^th^ percentile of the NME distribution of dormant genes in Group3 and, for cells expressing the active phenotype (aHSCs), the 99^th^ percentile of the NME distribution of active genes in Group1. These thresholds defined in normal homeostatic conditions, where then applied to all samples. Cells with NME values of dormant and active genes below the respective thresholds were considered as cells transiting among the two phenotypic states (trHSCs).

### Gene signatures

The WikiPathways lists of genes for Mus Musculus were downloaded using the Virtuoso SPARQL Query Editor for WikiPathways at http://sparql.wikipathways.org/.

We manually curated two gene signatures for dHSCs (Do28) and for aHSCs (Act166) considering both the bulk RNAseq and the single cell RNAseq raw data available in Cabezas *et al.* (43). In particular, we performed GSEA of the bulk RNAseq dataset using as query the list of genes upregulated in dormant or active HSCs at single cell level, respectively. Leading Edge genes of the enrichment profile represent our dormant (Do28; n=28) or active (Act166; n=166) HSC gene set and are listed in table S2A.

### Survival and differential expression analysis on TCGA AML patients

Clinical, cytogenetic, mutational and RNA-seq data from the TCGA-LAML cohort were downloaded from cBioPortal database (http://www.cbioportal.org/)(65) and 173 out of 200 patients were selected based on availability of gene expression data. RSEM expression values were pre-processed with nonparanormal transformation using the R package huge(49, 66) . Patients were reclassified into risk classes according to European Leukemia Net guidelines(49) based on their cytogenetic and mutational profile; FLT3-ITD mutant patients were always considered at intermediate risk.

To investigate the impact of TGFB1 expression on survival and relapse, we performed ROC analysis with the R package pROC(67) censoring patients at 17.1 months (the median overall survival of the analyzed cohort) and using TGFB1 expression as predictor. Best threshold for overall survival prediction was derived according to Youden index method and used to divide patients into high and low TGFB1 expression. For the two groups, overall survival probability was estimated by Kaplan-Meier method and compared by Logrank test, while cumulative incidence of relapse was tested by Gray’s method. For multivariate models, Cox Proportional Hazard Models were performed including total number of white blood cells, age, ELN risk class and TGFB1 status at diagnosis as covariates and using death by any cause and relapse as competing risks. Survival analyses were performed using R packages survminer and cmprsk. Clinical and molecular characteristics of patients in the two groups were compared by Pearson’s Chi-squared test for categorical variables and Kruskal-Wallis rank sum test for numerical variables. To examine differentially expressed genes in TGFB1 high vs low expressors, RNA-seq raw read counts were downloaded from the NCI Genomic Data Commons (https://gdc.cancer.gov/)(68), matched by barcode and then analyzed with the R package DESeq2(69) at gene level. The resulting DEGs list (adjusted p-value (q-value) < 0.1) was used to perform GSEAs.

## Acknowledgments

We thank L. Rotta and the Sequencing Facility at the IEO Genomic Unit. We thank C. Savino for technical assistance and Stefania Averaimo for critical reading of the manuscript. We thank Luca Mazzarella, Fernando Palluzzi and Emanuele Bonetti for bioinformatic advices.

## Author Contributions

MEBM, MM, CR and AP conducted the experiments. LL, CC and RH performed the bioinformatic analyses. GDC contributed to *in vivo* experiments. MF performed the imaging analyses. VT, AC and SP performed the histopathologic analysis. EC and PGP designed the experiments, oversaw the study and wrote the paper.

## Competing Interests

The authors declare they have no competing interests

All animal procedures were performed in accordance with the Italian Legislation.

## References

1. Welch JS, Ley TJ, Link DC, Miller CA, Larson DE, Koboldt DC, et al. The origin and evolution of mutations in acute myeloid leukemia. Cell. 2012;150(2):264–78.

2. Papaemmanuil E, Gerstung M, Bullinger L, Gaidzik VI, Paschka P, Roberts ND, et al. Genomic Classification and Prognosis in Acute Myeloid Leukemia. The New England journal of medicine. 2016;374(23):2209–21.

3. Mallardo M, Caronno A, Pruneri G, Raviele PR, Viale A, Pelicci PG, et al. NPMc+ and FLT3_ITD mutations cooperate in inducing acute leukaemia in a novel mouse model. Leukemia. 2013;27(11):2248–51.

4. Vassiliou GS, Cooper JL, Rad R, Li J, Rice S, Uren A, et al. Mutant nucleophosmin and cooperating pathways drive leukemia initiation and progression in mice. Nat Genet. 2011;43(5):470–5.

5. Mupo A, Celani L, Dovey O, Cooper JL, Grove C, Rad R, et al. A powerful molecular synergy between mutant Nucleophosmin and Flt3-ITD drives acute myeloid leukemia in mice. Leukemia. 2013;27(9):1917–20.

6. Loberg MA, Bell RK, Goodwin LO, Eudy E, Miles LA, SanMiguel JM, et al. Sequentially inducible mouse models reveal that Npm1 mutation causes malignant transformation of Dnmt3a- mutant clonal hematopoiesis. Leukemia. 2019;33(7):1635–49.

7. Guryanova OA, Shank K, Spitzer B, Luciani L, Koche RP, Garrett-Bakelman FE, et al. DNMT3A mutations promote anthracycline resistance in acute myeloid leukemia via impaired nucleosome remodeling. Nature medicine. 2016;22(12):1488–95.

8. Papaemmanuil E, Gerstung M, Malcovati L, Tauro S, Gundem G, Van Loo P, et al. Clinical and biological implications of driver mutations in myelodysplastic syndromes. Blood. 2013;122(22):3616–27; quiz 99.

9. Bowman RL, Busque L, Levine RL. Clonal Hematopoiesis and Evolution to Hematopoietic Malignancies. Cell Stem Cell. 2018;22(2):157–70.

10. Shlush LI, Zandi S, Mitchell A, Chen WC, Brandwein JM, Gupta V, et al. Identification of pre-leukaemic haematopoietic stem cells in acute leukaemia. Nature. 2014;506(7488):328–33.

11. Uckelmann HJ, Kim SM, Wong EM, Hatton C, Giovinazzo H, Gadrey JY, et al. Therapeutic targeting of preleukemia cells in a mouse model of NPM1 mutant acute myeloid leukemia. Science. 2020;367(6477):586–90.

12. Thol F, Klesse S, Kohler L, Gabdoulline R, Kloos A, Liebich A, et al. Acute myeloid leukemia derived from lympho-myeloid clonal hematopoiesis. Leukemia. 2017;31(6):1286–95.

13. Miles LA, Bowman RL, Merlinsky TR, Csete IS, Ooi AT, Durruthy-Durruthy R, et al. Single-cell mutation analysis of clonal evolution in myeloid malignancies. Nature. 2020;587(7834):477–82.

14. Kronke J, Bullinger L, Teleanu V, Tschurtz F, Gaidzik VI, Kuhn MW, et al. Clonal evolution in relapsed NPM1-mutated acute myeloid leukemia. Blood. 2013;122(1):100–8.

15. Mer AS, Heath EM, Madani Tonekaboni SA, Dogan-Artun N, Nair SK, Murison A, et al. Biological and therapeutic implications of a unique subtype of NPM1 mutated AML. Nat Commun. 2021;12(1):1054.

16. Martelli MP, Pettirossi V, Thiede C, Bonifacio E, Mezzasoma F, Cecchini D, et al. CD34+ cells from AML with mutated NPM1 harbor cytoplasmic mutated nucleophosmin and generate leukemia in immunocompromised mice. Blood. 2010;116(19):3907–22.

17. Pasqualucci L, Liso A, Martelli MP, Bolli N, Pacini R, Tabarrini A, et al. Mutated nucleophosmin detects clonal multilineage involvement in acute myeloid leukemia: Impact on WHO classification. Blood. 2006;108(13):4146–55.

18. Jan M, Snyder TM, Corces-Zimmerman MR, Vyas P, Weissman IL, Quake SR, et al. Clonal evolution of preleukemic hematopoietic stem cells precedes human acute myeloid leukemia. Sci Transl Med. 2012;4(149):149ra18.

19. Zeisig BB, Fung TK, Zarowiecki M, Tsai CT, Luo H, Stanojevic B, et al. Functional reconstruction of human AML reveals stem cell origin and vulnerability of treatment-resistant MLL-rearranged leukemia. Sci Transl Med. 2021;13(582).

20. Yun H, Narayan N, Vohra S, Giotopoulos G, Mupo A, Madrigal P, et al. Mutational synergy during leukemia induction remodels chromatin accessibility, histone modifications and three- dimensional DNA topology to alter gene expression. Nat Genet. 2021;53(10):1443–55.

21. Isobe T, Kucinski I, Barile M, Wang X, Hannah R, Bastos HP, et al. Preleukemic single-cell landscapes reveal mutation-specific mechanisms and gene programs predictive of AML patient outcomes. Cell Genom. 2023;3(12):100426.

22. Srinivas S, Watanabe T, Lin CS, William CM, Tanabe Y, Jessell TM, et al. Cre reporter strains produced by targeted insertion of EYFP and ECFP into the ROSA26 locus. BMC developmental biology. 2001;1:4.

23. Morrison SJ, Weissman IL. The long-term repopulating subset of hematopoietic stem cells is deterministic and isolatable by phenotype. Immunity. 1994;1(8):661–73.

24. Kent DG, Copley MR, Benz C, Wohrer S, Dykstra BJ, Ma E, et al. Prospective isolation and molecular characterization of hematopoietic stem cells with durable self-renewal potential. Blood. 2009;113(25):6342–50.

25. Dovey OM, Cooper JL, Mupo A, Grove CS, Lynn C, Conte N, et al. Molecular synergy underlies the co-occurrence patterns and phenotype of NPM1-mutant acute myeloid leukemia. Blood. 2017;130(17):1911–22.

26. Wilson A, Laurenti E, Oser G, van der Wath RC, Blanco-Bose W, Jaworski M, et al. Hematopoietic stem cells reversibly switch from dormancy to self-renewal during homeostasis and repair. Cell. 2008;135(6):1118–29.

27. Chen CL, Faltusova K, Molik M, Savvulidi F, Chang KT, Necas E. Low c-Kit Expression Level Induced by Stem Cell Factor Does Not Compromise Transplantation of Hematopoietic Stem Cells. Biol Blood Marrow Transplant. 2016;22(7):1167–72.

28. Feil R, Brocard J, Mascrez B, LeMeur M, Metzger D, Chambon P. Ligand-activated site- specific recombination in mice. Proceedings of the National Academy of Sciences of the United States of America. 1996;93(20):10887–90.

29. Challen GA, Sun D, Jeong M, Luo M, Jelinek J, Berg JS, et al. Dnmt3a is essential for hematopoietic stem cell differentiation. Nature genetics. 2011;44(1):23–31.

30. Hu Y, Smyth GK. ELDA: extreme limiting dilution analysis for comparing depleted and enriched populations in stem cell and other assays. J Immunol Methods. 2009;347(1-2):70–8.

31. Zou P, Yoshihara H, Hosokawa K, Tai I, Shinmyozu K, Tsukahara F, et al. p57(Kip2) and p27(Kip1) cooperate to maintain hematopoietic stem cell quiescence through interactions with Hsc70. Cell stem cell. 2011;9(3):247–61.

32. Essers MA, Offner S, Blanco-Bose WE, Waibler Z, Kalinke U, Duchosal MA, et al. IFNalpha activates dormant haematopoietic stem cells in vivo. Nature. 2009;458(7240):904–8.

33. Lerner C, Harrison DE. 5-Fluorouracil spares hemopoietic stem cells responsible for long-term repopulation. Experimental hematology. 1990;18(2):114–8.

34. Chu SH, Heiser D, Li L, Kaplan I, Collector M, Huso D, et al. FLT3-ITD knockin impairs hematopoietic stem cell quiescence/homeostasis, leading to myeloproliferative neoplasm. Cell stem cell. 2012;11(3):346–58.

35. Baldridge MT, King KY, Boles NC, Weksberg DC, Goodell MA. Quiescent haematopoietic stem cells are activated by IFN-gamma in response to chronic infection. Nature. 2010;465(7299):793–7.

36. Bernitz JM, Kim HS, MacArthur B, Sieburg H, Moore K. Hematopoietic Stem Cells Count and Remember Self-Renewal Divisions. Cell. 2016;167(5):1296–309 e10.

37. Kuhn R, Schwenk F, Aguet M, Rajewsky K. Inducible gene targeting in mice. Science. 1995;269(5229):1427–9.

38. Lee BH, Tothova Z, Levine RL, Anderson K, Buza-Vidas N, Cullen DE, et al. FLT3 mutations confer enhanced proliferation and survival properties to multipotent progenitors in a murine model of chronic myelomonocytic leukemia. Cancer Cell. 2007;12(4):367–80.

39. Alcalay M, Tiacci E, Bergomas R, Bigerna B, Venturini E, Minardi SP, et al. Acute myeloid leukemia bearing cytoplasmic nucleophosmin (NPMc+ AML) shows a distinct gene expression profile characterized by up-regulation of genes involved in stem-cell maintenance. Blood. 2005;106(3):899–902.

40. Brunetti L, Gundry MC, Sorcini D, Guzman AG, Huang YH, Ramabadran R, et al. Mutant NPM1 Maintains the Leukemic State through HOX Expression. Cancer Cell. 2018;34(3):499–512 e9.

41. Venezia TA, Merchant AA, Ramos CA, Whitehouse NL, Young AS, Shaw CA, et al. Molecular signatures of proliferation and quiescence in hematopoietic stem cells. PLoS biology. 2004;2(10):e301.

42. Yamada T, Park CS, Lacorazza HD. Genetic control of quiescence in hematopoietic stem cells. Cell cycle. 2013;12(15):2376–83.

43. Cabezas-Wallscheid N, Buettner F, Sommerkamp P, Klimmeck D, Ladel L, Thalheimer FB, et al. Vitamin A-Retinoic Acid Signaling Regulates Hematopoietic Stem Cell Dormancy. Cell. 2017;169(5):807–23 e19.

44. Haas S, Trumpp A, Milsom MD. Causes and Consequences of Hematopoietic Stem Cell Heterogeneity. Cell Stem Cell. 2018;22(5):627–38.

45. Wilson NK, Kent DG, Buettner F, Shehata M, Macaulay IC, Calero-Nieto FJ, et al. Combined Single-Cell Functional and Gene Expression Analysis Resolves Heterogeneity within Stem Cell Populations. Cell Stem Cell. 2015;16(6):712–24.

46. Naka K, Hoshii T, Muraguchi T, Tadokoro Y, Ooshio T, Kondo Y, et al. TGF-beta-FOXO signalling maintains leukaemia-initiating cells in chronic myeloid leukaemia. Nature. 2010;463(7281):676–80.

47. Prunier C, Baker D, Ten Dijke P, Ritsma L. TGF-beta Family Signaling Pathways in Cellular Dormancy. Trends Cancer. 2019;5(1):66–78.

48. Cancer Genome Atlas Research N, Ley TJ, Miller C, Ding L, Raphael BJ, Mungall AJ, et al. Genomic and epigenomic landscapes of adult de novo acute myeloid leukemia. N Engl J Med. 2013;368(22):2059–74.

49. Dohner H, Estey E, Grimwade D, Amadori S, Appelbaum FR, Buchner T, et al. Diagnosis and management of AML in adults: 2017 ELN recommendations from an international expert panel. Blood. 2017;129(4):424–47.

50. Montalban-Bravo G, Kanagal-Shamanna R, Sasaki K, Patel K, Ganan-Gomez I, Jabbour E, et al. NPM1 mutations define a specific subgroup of MDS and MDS/MPN patients with favorable outcomes with intensive chemotherapy. Blood Adv. 2019;3(6):922–33.

51. Krivtsov AV, Figueroa ME, Sinha AU, Stubbs MC, Feng Z, Valk PJ, et al. Cell of origin determines clinically relevant subtypes of MLL-rearranged AML. Leukemia. 2013;27(4):852–60.

52. Cozzio A, Passegue E, Ayton PM, Karsunky H, Cleary ML, Weissman IL. Similar MLL-associated leukemias arising from self-renewing stem cells and short-lived myeloid progenitors. Genes Dev. 2003;17(24):3029–35.

53. So CW, Karsunky H, Passegue E, Cozzio A, Weissman IL, Cleary ML. MLL-GAS7 transforms multipotent hematopoietic progenitors and induces mixed lineage leukemias in mice. Cancer Cell. 2003;3(2):161–71.

54. Ayton PM, Cleary ML. Transformation of myeloid progenitors by MLL oncoproteins is dependent on Hoxa7 and Hoxa9. Genes Dev. 2003;17(18):2298–307.

55. Krivtsov AV, Evans K, Gadrey JY, Eschle BK, Hatton C, Uckelmann HJ, et al. A Menin-MLL Inhibitor Induces Specific Chromatin Changes and Eradicates Disease in Models of MLL- Rearranged Leukemia. Cancer Cell. 2019;36(6):660–73 e11.

56. Carraway HE. Treatment options for patients with myelodysplastic syndromes after hypomethylating agent failure. Hematology Am Soc Hematol Educ Program. 2016;2016(1):470–7.

57. Gil-Perez A, Montalban-Bravo G. Management of myelodysplastic syndromes after failure of response to hypomethylating agents. Ther Adv Hematol. 2019;10:2040620719847059.

58. Houshmand M, Simonetti G, Circosta P, Gaidano V, Cignetti A, Martinelli G, et al. Chronic myeloid leukemia stem cells. Leukemia. 2019;33(7):1543–56.

59. Li Q, Bohin N, Wen T, Ng V, Magee J, Chen SC, et al. Oncogenic Nras has bimodal effects on stem cells that sustainably increase competitiveness. Nature. 2013;504(7478):143-7.

60. Furia L, Pelicci PG, Faretta M. A computational platform for robotized fluorescence microscopy (II): DNA damage, replication, checkpoint activation, and cell cycle progression by high-content high-resolution multiparameter image-cytometry. Cytometry A. 2013;83(4):344–55.

61. Kim D, Pertea G, Trapnell C, Pimentel H, Kelley R, Salzberg SL. TopHat2: accurate alignment of transcriptomes in the presence of insertions, deletions and gene fusions. Genome Biol. 2013;14(4):R36.

62. Anders S, Pyl PT, Huber W. HTSeq--a Python framework to work with high-throughput sequencing data. Bioinformatics. 2015;31(2):166–9.

63. Robinson MD, McCarthy DJ, Smyth GK. edgeR: a Bioconductor package for differential expression analysis of digital gene expression data. Bioinformatics. 2010;26(1):139–40.

64. Qiu X, Mao Q, Tang Y, Wang L, Chawla R, Pliner HA, et al. Reversed graph embedding resolves complex single-cell trajectories. Nat Methods. 2017;14(10):979–82.

65. Cerami E, Gao J, Dogrusoz U, Gross BE, Sumer SO, Aksoy BA, et al. The cBio cancer genomics portal: an open platform for exploring multidimensional cancer genomics data. Cancer Discov. 2012;2(5):401–4.

66. Zhao T, Liu H, Roeder K, Lafferty J, Wasserman L. The huge Package for High- dimensional Undirected Graph Estimation in R. J Mach Learn Res. 2012;13:1059–62.

67. Robin X, Turck N, Hainard A, Tiberti N, Lisacek F, Sanchez JC, et al. pROC: an open- source package for R and S+ to analyze and compare ROC curves. BMC Bioinformatics. 2011;12:77.

68. Grossman RL, Heath AP, Ferretti V, Varmus HE, Lowy DR, Kibbe WA, et al. Toward a Shared Vision for Cancer Genomic Data. N Engl J Med. 2016;375(12):1109–12.

69. Love MI, Huber W, Anders S. Moderated estimation of fold change and dispersion for RNA-seq data with DESeq2. Genome Biol. 2014;15(12):550.

70. Billing M, Rorby E, May G, Tipping AJ, Soneji S, Brown J, et al. A network including TGFbeta/Smad4, Gata2, and p57 regulates proliferation of mouse hematopoietic progenitor cells. Exp Hematol. 2016;44(5):399–409 e5.

