## Supplemental Figures 1-5 for "NPMc+ and Flt3-ITD cooperation promotes a new oncogenic HSC state that supports transformation and leukemia stem cell maintenance"

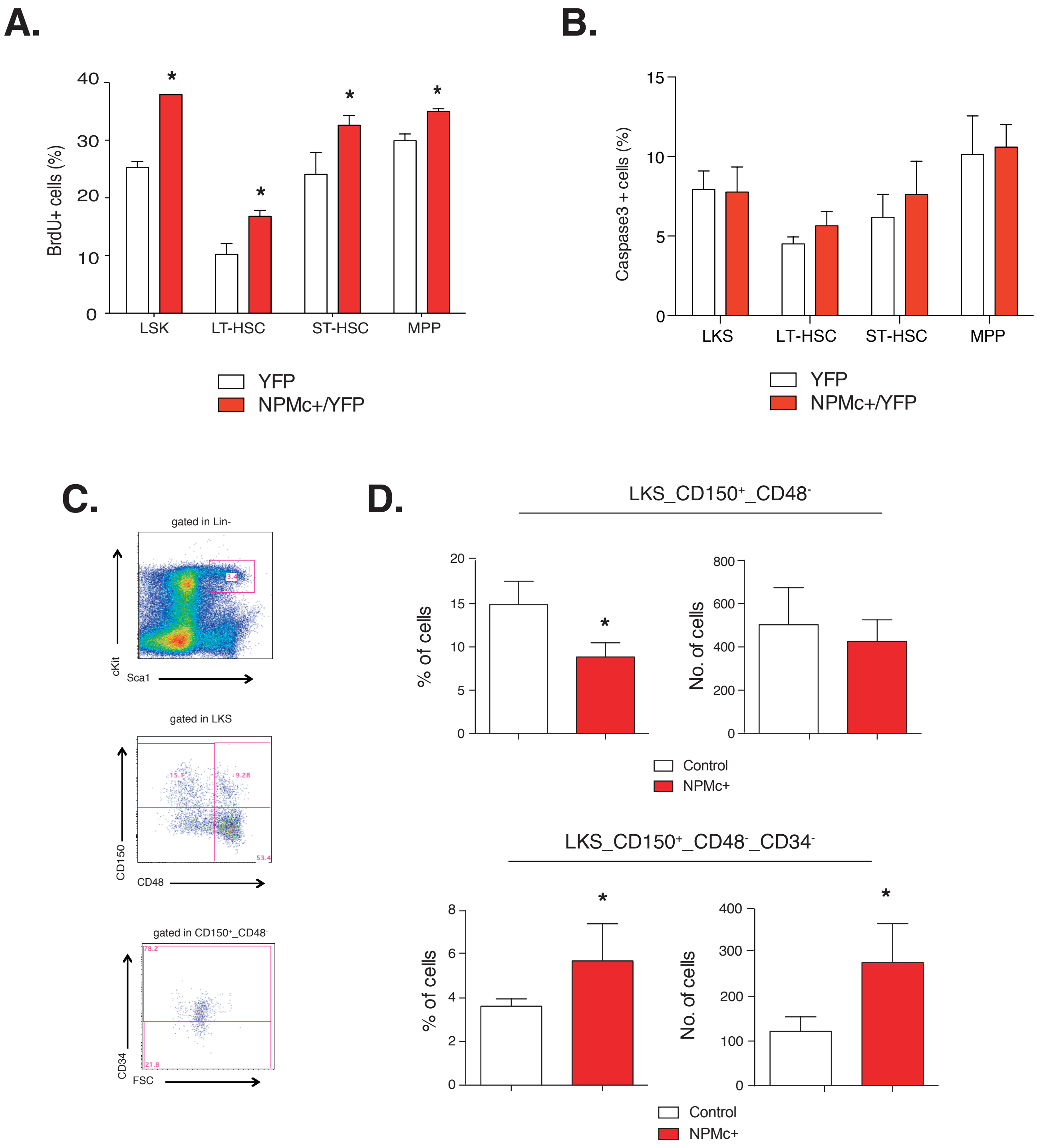

A.
BrdU+ cells (%)
LSK
LT-HSC
ST-HSC
MPP
YFP
NPMc+/YFP
B.
Caspase3 + cells (%)
LKS
LT-HSC
ST-HSC
MPP
YFP
NPMc+/YFP
C.
gated in Lin-
cKit
Sca1
gated in LKS
CD150
CD48
gated in CD150+_CD48-
CD34
FSC
D.
LKS_CD150+_CD48-
% of cells
No. of cells
Control
NPMc+
LKS_CD150+_CD48-_CD34-
% of cells
No. of cells
Control
NPMc+


**Figure S1**

**Figure S1. Analyses of different HSPCs population upon NPMc+ expression. A.** *In vivo* BrdU assay: BM was collected 12 hours after *in vivo* BrdU administration and analysed by FACS to evaluate the percentage of BrdU positive cells in different HSPCs population (LSKs, LT-HSCs, ST-HSCs, MPPs) (3-5 mice per cohort; graph representative of 1 of 2 independent experiments; Unpaired Student's t test has been applied, *=$p<0.05$)**. B.** Percentage of Caspase-3 positive cells in the indicated HSPC subpopulations (3 mice per cohort). Mean ± s.d. values are shown. **C.** Representative FACS gating schemes for the analysis of different BM populations. Upper panel: gating of the LSK (c-Kit+, Sca-1+, Lin-) population within lineage negative cells; middle panel: gating of CD48-/CD150+ population within the LSK cells; lower panel: gating of the CD34- population within the CD48-/CD150+ cells. **D.** Left panels: percentages of LSK/CD48-/CD150+ (upper) and LSK/CD48-/CD150+/CD34- BM subpopulations as gated in C. Right panels: numbers (per million of BM-MNCs) of the same populations (3-5 mice per cohort; graph representative of 1 of 2 independent experiments; *=$p<0.05$).

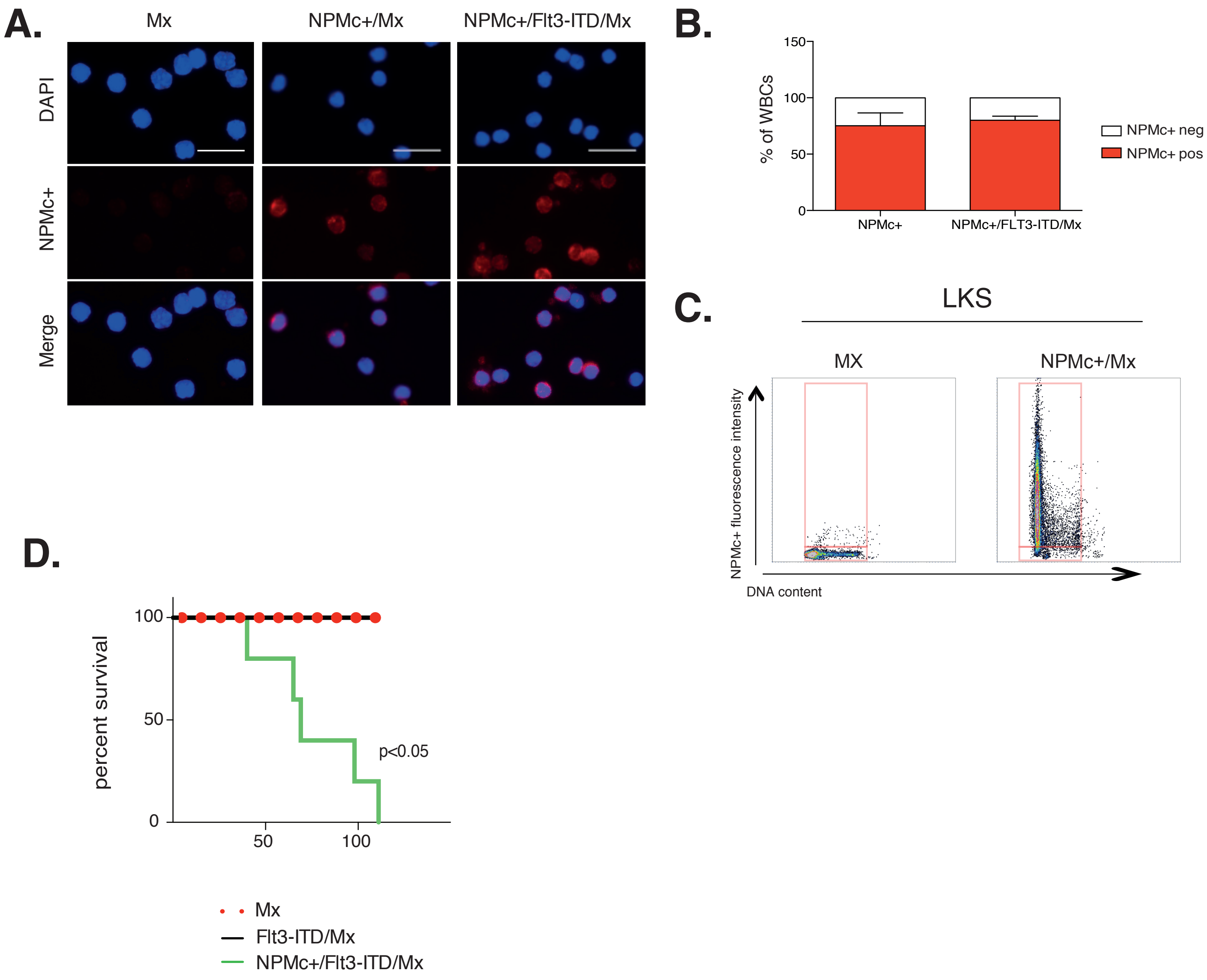
A.
Mx
NPMc+/Mx
NPMc+/Flt3-ITD/Mx
DAPI
NPMc+
Merge
B.
150
100
50
0
% of WBCs
NPMc+
NPMc+/FLT3-ITD/Mx
NPMc+ neg
NPMc+ pos
C.
LKS
MX
NPMc+/Mx
NPMc+ fluorescence intensity
DNA content
D.
100
50
0
percent survival
50
100
p<0.05
Mx
Flt3-ITD/Mx
NPMc+/Flt3-ITD/Mx

Figure S2

**Figure S2. Mx-CRE mediated NPMc+ expression in the PB and in the LKS compartment**

**A.** Representative images of the immunofluorescence analysis performed on Cytospin® preparations of PB WBCs derived from pIpC treated Mx, NPMc+ and NPMc+/FLT3-ITD mice. In red, rabbit polyclonal antibody against NPMc+; in blue, DAPI staining of nuclei. NPMc+ cytoplasmic localization is confirmed by merging red and blue staining. Scale bar 100 um. **B**. Histograms represent the percentage of NPMc+ positive WBCs in the total WBCs population (data represent the pool of four independent experiments). **C**. Plots represent the NPMc+ dependent fluorescence signal intensity/cell, quantified by the A.M.I.CO. computational platform [1] in Mx and NPMc+/Mx LKS samples. DNA content was evaluated using DAPI staining intensity (data represent the pool of two independent experiments). **D.** Kaplan Meyer survival curve for Mx, Flt3-ITD/Mx and NPMc+/Flt3-ITD/Mx mice (5 animals per group, p-value calculated using the log-rank test).

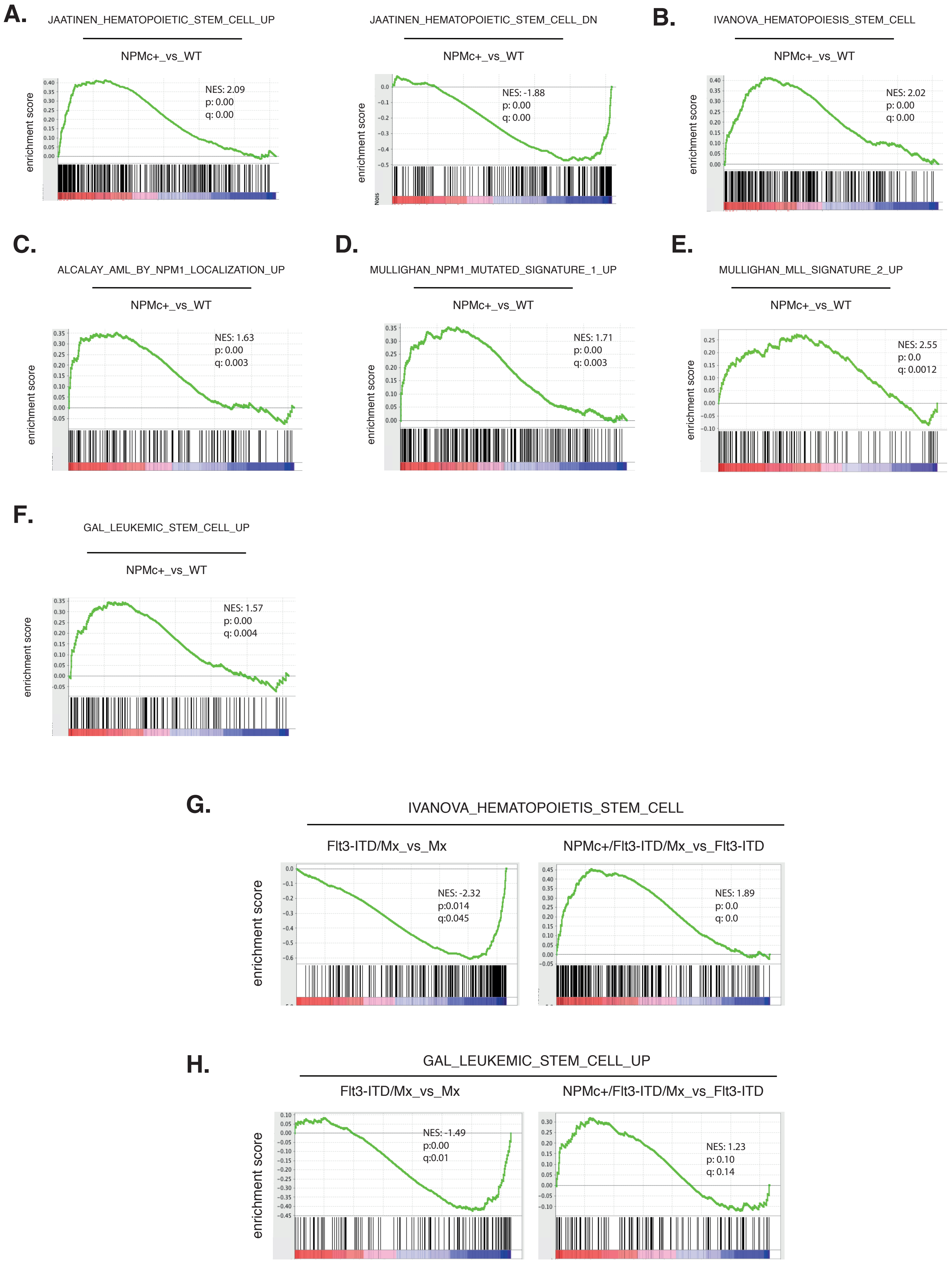
A.
JAATINEN_HEMATOPOIETIC_STEM_CELL_UP
NPMc+_vs_WT
enrichment score
NES: 2.09
p: 0.00
q: 0.00
JAATINEN_HEMATOPOIETIC_STEM_CELL_DN
NPMc+_vs_WT
enrichment score
NES: -1.88
p: 0.00
q: 0.00
B.
IVANOVA_HEMATOPOIESIS_STEM_CELL
NPMc+_vs_WT
enrichment score
NES: 2.02
p: 0.00
q: 0.00
C.
ALCALAY_AML_BY_NPM1_LOCALIZATION_UP
NPMc+_vs_WT
enrichment score
NES: 1.63
p: 0.00
q: 0.003
D.
MULLIGHAN_NPM1_MUTATED_SIGNATURE_1_UP
NPMc+_vs_WT
enrichment score
NES: 1.71
p: 0.00
q: 0.003
E.
MULLIGHAN_MLL_SIGNATURE_2_UP
NPMc+_vs_WT
enrichment score
NES: 2.55
p: 0.0
q: 0.0012
F.
GAL_LEUKEMIC_STEM_CELL_UP
NPMc+_vs_WT
enrichment score
NES: 1.57
p: 0.00
q: 0.004
G.
IVANOVA_HEMATOPOIETIS_STEM_CELL
Flt3-ITD/Mx_vs_Mx
NPMc+/Flt3-ITD/Mx_vs_Flt3-ITD
enrichment score
NES: -2.32
p:0.014
q:0.045
NES: 1.89
p: 0.0
q: 0.0
H.
GAL_LEUKEMIC_STEM_CELL_UP
Flt3-ITD/Mx_vs_Mx
NPMc+/Flt3-ITD/Mx_vs_Flt3-ITD
enrichment score
NES: -1.49
p:0.00
q:0.01
NES: 1.23
p: 0.10
q: 0.14

**Figure S3**

**Figure S3. NPMc+ induces the expression of HSC and LSC related signatures**

**A-F**. Gene expression microarray data were used to identify gene sets enriched in NPMc+/YFP LT-HSCs compared to YFP LT-HSCs. GSEAs show a significant correlation of the NPMc+ LT-HSC gene expression profile with: i) genes upregulated (**A** left panel) and downregulated (**A** right panel and **B**) in human hematopoietic stem cells[2,3]; ii) genes upregulated in NPMc+[4] and MLL[5] human AMLs (**C**; **D-E**); iii) genes up regulated in AML stem cells[6] (**F**). **G-H**. RNAseq data were used to identify enriched gene sets in Flt3-ITD/Mx vs Mx LT-HSCs (left panels) or NPMc+/Flt3-ITD/Mx vs Flt3-ITD/Mx LT-HSCs (right panels). GSEAs show significant correlations with genes upregulated in human hematopoietic stem cells (**G**), and genes upregulated in AML stem cells (**H**). Normalized enrichment score (NES), p value (p) and false discovery rate (q) are indicated in each panel.

**A.**

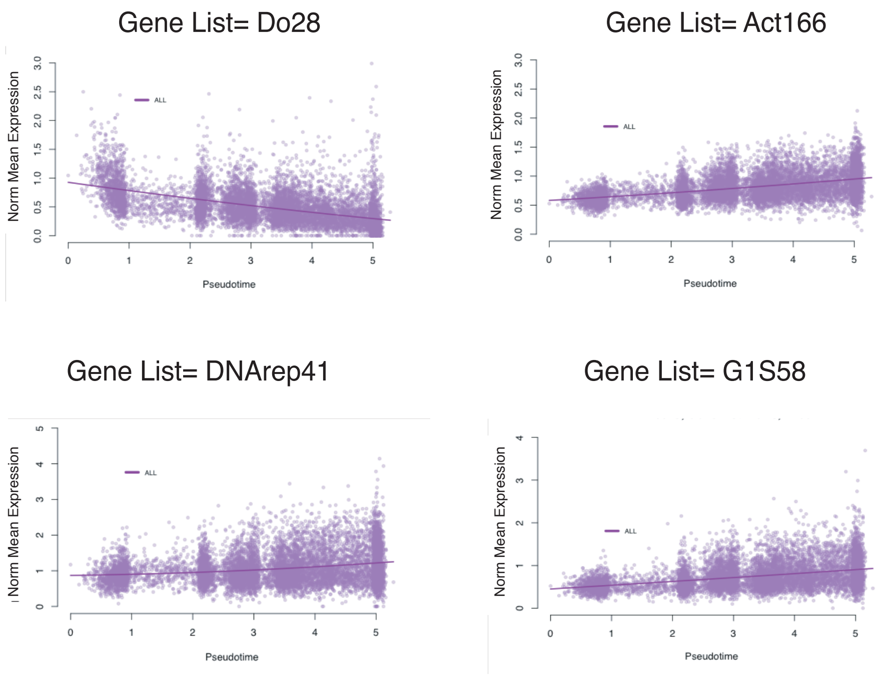

Gene List= Do28
Gene List= Act166
Gene List= DNArep41
Gene List= G1S58
Norm Mean Expression
Pseudotime
ALL


**B.**

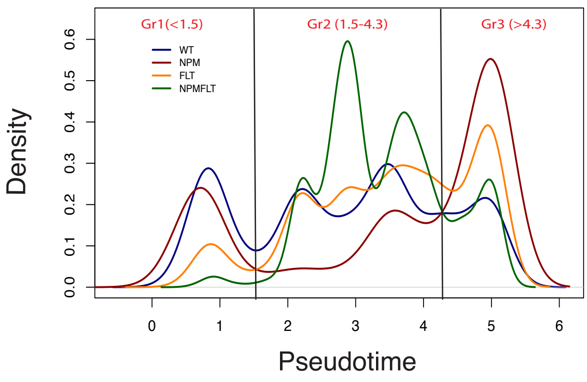

Gr1(<1.5)
Gr2 (1.5-4.3)
Gr3 (>4.3)
WT
NPM
FLT
NPMFLT
Density
Pseudotime


Figure S4

**Figure S4. Single cell RNAseq analysis**. **A**. scatter plots showing normalized mean expression levels by pseudotime of gene lists reported in Table S2, in the pool of the four samples. Do28 (upper left panel, dHSC related signature); Act166 (upper right panel, aHSC related signature); DNArep41 (lower left panel, genes annotated to the DNA replication pathway from WikiPathways[7]); G1S58 (lower right panel, genes annotated to G1 to S cell cycle transition from WikiPathways). Colored lines depict the polynomial fit of the normalized expression values. **C.** density plot displaying single cell distribution along the trajectory. Groups1-3 were defined as described in materials and methods section (pseudotime <1.5; > 1.5 and <4.3; >4.3, respectively).

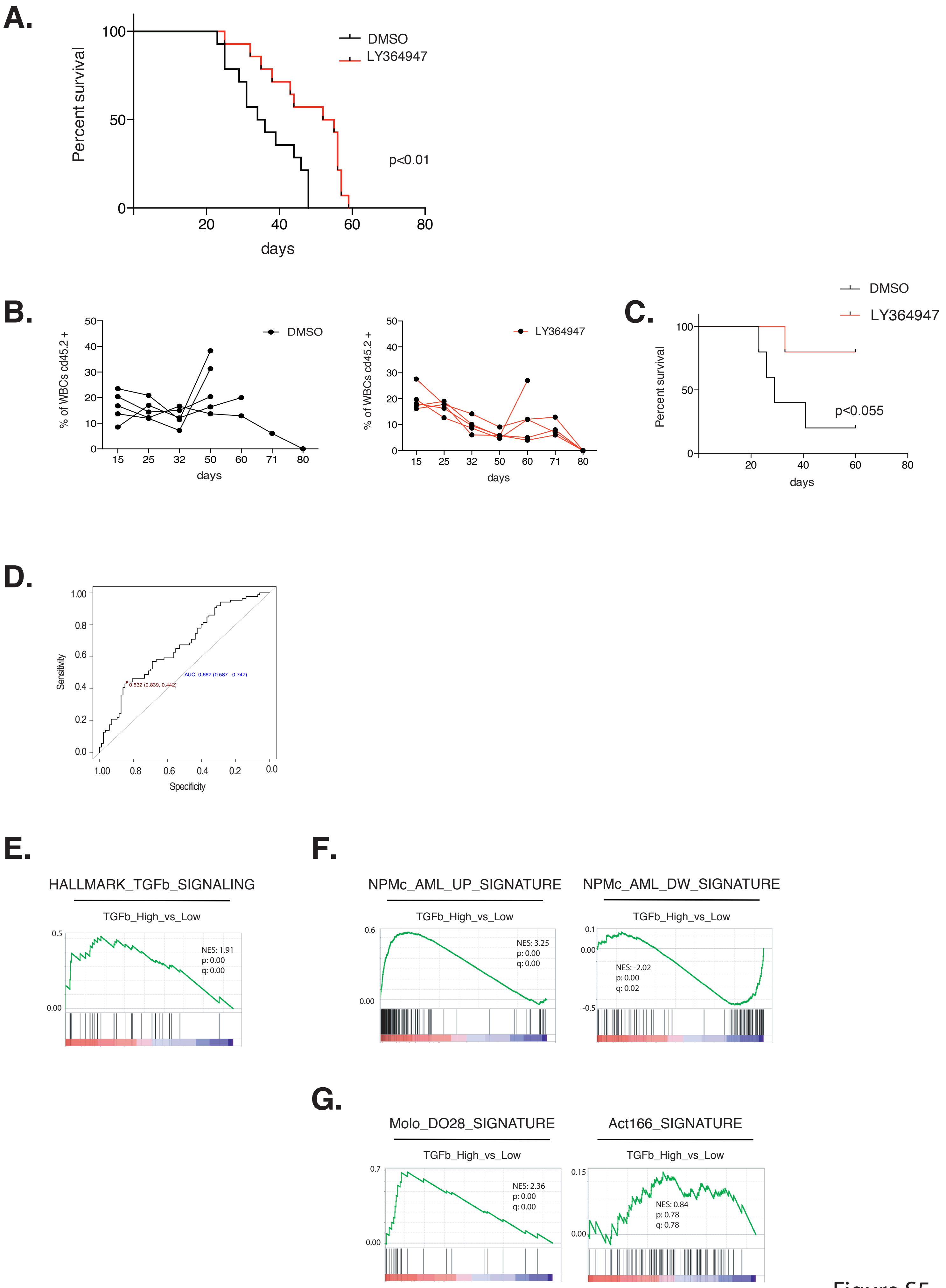
A.
Percent survival
100
50
0
20
40
60
80
days
DMSO
LY364947
p<0.01
B.
% of WBCs cd45.2 +
50
40
30
20
10
0
15
25
32
50
60
71
80
days
DMSO
LY364947
C.
DMSO
LY364947
Percent survival
100
50
0
20
40
60
80
days
p<0.055
D.
Sensitivity
1.00
0.8
0.6
0.4
0.2
0.0
Specificity
0.532 (0.839, 0.442)
AUC: 0.667 (0.587...0.747)
E.
HALLMARK_TGFb_SIGNALING
TGFb_High_vs_Low
0.5
0.00
NES: 1.91
p: 0.00
q: 0.00
F.
NPMc_AML_UP_SIGNATURE
TGFb_High_vs_Low
0.6
0.00
NES: 3.25
p: 0.00
q: 0.00
NPMc_AML_DW_SIGNATURE
TGFb_High_vs_Low
0.1
0.00
-0.5
NES: -2.02
p: 0.00
q: 0.02
G.
Molo_DO28_SIGNATURE
TGFb_High_vs_Low
0.7
0.00
NES: 2.36
p: 0.00
q: 0.00
Act166_SIGNATURE
TGFb_High_vs_Low
0.15
0.00
NES: 0.84
p: 0.78
q: 0.78

Figure S5

**Figure S5. Pharmacological inhibition of TGFβ pathway affects AML growth *in vivo*. A**. Kaplan Meyer survival curve of mice transplanted with a NPMc+/*Flt3-ITD* murine AML and treated with the LY364947 TGFβR-I inhibitor or vehicle every other day for 10 days (10 animals/group). **B**. Graphs represent the percentage of control (left panel) or TGFβRI inhibited (right panel) blasts (CD45.2+) in the PB of secondary transplanted mice (n=5 animals per group) **C.** Kaplan Meier survival curve of mice transplanted with TGFβRI inhibited blasts or control blasts (n=5 animals). The p value was calculated with the log rank test. **D.** ROC analysis to identify best *TGFβ1* expression cutoff-predictor by Youden index (red); AUC: area under the curve (blue). **E-G.** RNAseq data from patients (TCGA data set) were used to identify enriched gene signatures in TGFβ1 high versus TGFβ1 low groups. GSEAs show a significant correlation with: i) HALLMARK TGFβ pathway genes; ii) genes upregulated (**F** left panel) and downregulated (**F** right panel) in NPMc+[4] human AMLs ; iii) HSCs dormant (Do28+MolO; Table S2A)) gene signatures (**G** left panel). No correlation was found with the HSCs active (Act166; Table S2A) gene signature (**G** right panel). Normalized enrichment score (NES), p value (p) and false discovery rate (q) are indicated in each panel.

Supplementary References


1 Furia, L., Pelicci, P. G. & Faretta, M. A computational platform for robotized fluorescence microscopy (II): DNA damage, replication, checkpoint activation, and cell cycle progression by high-content high-resolution multiparameter image-cytometry. *Cytometry A* 83, 344-355, doi:10.1002/cyto.a.22265 (2013).

2 Jaatinen, T. *et al.* Global gene expression profile of human cord blood-derived CD133+ cells. *Stem Cells* 24, 631-641, doi:10.1634/stemcells.2005-0185 (2006).

3 Ivanova, N. B. *et al.* A stem cell molecular signature. *Science* 298, 601-604, doi:10.1126/science.1073823 (2002).

4 Alcalay, M. *et al.* Acute myeloid leukemia bearing cytoplasmic nucleophosmin (NPMc+ AML) shows a distinct gene expression profile characterized by up-regulation of genes involved in stem-cell maintenance. *Blood* 106, 899-902, doi:2005-02-0560 [pii] 10.1182/blood-2005-02-0560 (2005).

5 Mullighan, C. G. *et al.* Pediatric acute myeloid leukemia with NPM1 mutations is characterized by a gene expression profile with dysregulated HOX gene expression distinct from MLL-rearranged leukemias. *Leukemia* 21, 2000-2009, doi:10.1038/sj.leu.2404808 (2007).

6 Gal, H. *et al.* Gene expression profiles of AML derived stem cells; similarity to hematopoietic stem cells. *Leukemia* 20, 2147-2154, doi:10.1038/sj.leu.2404401 (2006).

7 Kutmon, M. *et al.* WikiPathways: capturing the full diversity of pathway knowledge. *Nucleic Acids Res* 44, D488-494, doi:10.1093/nar/gkv1024 (2016).
